# The TWK-26/KCNK3 potassium channel and FLR-4 protein kinase coordinate nutrient absorption in the *C. elegans* intestine

**DOI:** 10.1101/2024.05.06.592787

**Authors:** Sarah K. Torzone, Trae Carroll, Peter C. Breen, Natalie R. Cohen, Kaylee N. Simmons, Kylie R. VanDerMolen, Keith W. Nehrke, Robert H. Dowen

## Abstract

Ion channels are necessary for proper water and nutrient absorption in the intestine, thereby supporting cellular metabolism and organismal growth. While a role for Na^+^ co-transporters and pumps in intestinal nutrient absorption is well defined, how individual K^+^ uniporters function in this process is poorly understood. Using *Caenorhabditis elegans*, we show that a gain-of-function mutation in *twk-26*, which encodes a two-pore domain K^+^ ion channel orthologous to human KCNK3, facilitates nutrient absorption and suppresses the metabolic and developmental defects caused by loss of DRL-1/FLR-4 signaling. Mutations in *drl-1* and *flr-4,* which encode two unique kinases that are components of a mitogen-activated protein kinase (MAPK) pathway, and the downstream *flr-1* Na^+^ ion channel, cause severe growth defects, reduced lipid storage, and a dramatic increase in autophagic lysosomes, which mirror dietary restriction phenotypes. We reveal that this dietary restriction phenotype is likely the result of impaired intestinal amino acid absorption, which is restored upon activation of TWK-26. Furthermore, we show that loss of *flr-4* disrupts intracellular and extracellular pH gradients, suggesting that the FLR-4 pathway may be necessary to maintain intestinal ion homeostasis and facilitate nutrient absorption. The altered pH gradients in the *flr-4* mutant are partially restored by activation of TWK-26, demonstrating a novel role for this K^+^ ion channel in governing intestinal physiology and metabolism.

## INTRODUCTION

Ion channels are membrane proteins that regulate the excitability of specialized cells that generate and propagate action potentials, as well as cell types that generate ion gradients to govern water, electrolyte, and small molecule transport across membranes. In the intestine, the maintenance of membrane potential and transmembrane ion gradients is essential for nutrient absorption and digestion (Lu et al. 2023). Dysregulation of ion channel expression or function in intestinal enterocytes contributes to several gastrointestinal (GI) pathologies, as seen in cystic fibrosis, colonic inflammation, and colorectal cancers (Anderson et al. 2019). For example, intermediate conductance potassium ion (IK) channels on the basolateral membrane of colonocytes are required for membrane hyperpolarization that facilitate Na^+^ and Cl^-^ transport across the cell membrane, which when impaired results in diseases such as ulcerative colitis (Al-Hazza et al. 2012). Consistently, decreased expression of basolateral IK channels in inflammatory bowel disease results in the secretion of K^+^ ions into the intestinal lumen, causing abdominal pain and diarrhea (Magalhães et al. 2016). Because ion channels are essential for preserving gastrointestinal function and organismal energy homeostasis, it is crucial to understand the molecular mechanisms that govern their activity in order to identify novel therapeutic targets of GI disease.

The depolarization or hyperpolarization of intestinal epithelial cells (IECs) is largely responsible for tuning the activity of nutrient transporters (Lu et al. 2023). For example, the transport of peptides by the di/tripeptide nutrient transporter PepT1 requires the recycling of an H^+^ ion gradient across the apical membrane of IECs (Thwaites et al. 2002). Thus, proper nutrient absorption in the intestine is pH dependent. The electrogenic forces that drive proton-coupled amino acid and glucose absorption in IECs are often stabilized by K^+^ channels (Lu et al. 2023), but how specific potassium ion channels regulate metabolic processes in the intestine to promote organismal homeostasis remains poorly understood.

Regulation of intestinal nutrient absorption by ion gradients is well conserved. In the nematode *Caenorhabditis elegans*, 20 epithelial cells absorb all the lipids, peptides, and carbohydrates required to maintain metabolic homeostasis and promote growth. The *C. elegans* gene *pept-1* encodes an intestinal oligopeptide transporter that is the ortholog of mammalian PepT1. As the primary di/tri-peptide transporter in the intestine, loss of *pept-1* impairs amino acid uptake, which slows development and reduces body size in *C. elegans* (Nehrke 2003). Likewise, loss of the apical Na^+^/H^+^ exchanger NHX-2/NHE3, an intestinal antiporter that maintains intestinal acidification and drives pH-dependent nutrient absorption, impairs growth and development (Nehrke 2003). Furthermore, disruption of *nhx-2* blunts lipid absorption, possibly arising from PEPT-1-dependent intracellular acidification (Spanier et al. 2009).

Intestinal pH oscillations, which support nutrient absorption and metabolic homeostasis, are also necessary to promote a rhythmic defecation cycle in *C. elegans*. The *C. elegans* defecation motor program (DMP) occurs every ∼45 seconds and is coordinated by calcium and proton oscillations in the intestine (Iwasaki et al. 1995; Branicky and Hekimi 2006; Allman et al. 2009; Pfeiffer et al. 2008). The DMP is coupled to the absorption of essential nutrients, including dipeptides, glucose, and lipids (Allman et al. 2009; Sheng et al. 2015; Suzuki et al. 2022). Consistently, mutations that disrupt the DMP (*e.g.*, *aex-5* and *flr-1*) reduce somatic intracellular lipid stores (Kaulich et al. 2022; Sheng et al. 2015), likely due to impaired intestinal acidification and poor nutrient absorption. It is known that cell autonomous calcium signaling drives the ultradian DMP, and the mechanisms that regulate intestinal calcium oscillations are well defined (Estevez and Strange 2005). Thus, maintenance of intestinal ion homeostasis is imperative to the metabolic homeostasis of the organism, governing both defecation and nutrient absorption; however, the molecular mechanisms that connect these ion-driven processes has remained elusive.

We previously demonstrated that DRL-1 and FLR-4, two intestinal protein kinases that function upstream of a mitogen-activated protein kinase (MAPK) signaling pathway, promote growth, lipid accumulation, and reproduction in *C. elegans* by, at least in part, suppressing the p38 innate immune pathway (Torzone et al. 2023). Loss-of-function alleles of *drl-1* (also known as *flr-3*) (Honey et al. 2023), *flr-4*, and *flr-1* were isolated in a forward genetic screen for mutations that confer resistance to fluoride (Katsura et al. 1994). The *flr-1* gene encodes a degenerin (DEG)/epithelial sodium channel (ENaC). Intriguingly, mutations in *drl-1*, *flr-4*, and *flr-1* all disrupt the DMP, cause developmental delay, and reduce lipid stores (Chamoli et al. 2014; Katsura et al. 1994; Kaulich et al. 2022; Take-uchi et al. 1998, 2005; Torzone et al. 2023), suggesting that DRL-1/FLR-4 signaling and ion channels may function in concert to coordinate defecation, metabolism, and development; however, the mechanistic basis of this regulation is unknown.

In this study, we use a forward genetic approach to isolate a gain-of-function mutation in *twk-26*, which encodes a potassium two-pore domain (K2P) ion channel, that suppresses the metabolic and developmental defects conferred by the *flr-4* and *flr-1* mutations. Additionally, we establish that an activating mutation that increases the open probability of TWK-26 mimics the gain-of-function *twk-26* mutation, suggesting that the role of TWK-26 in metabolic homeostasis is linked to its ion channel activity. Remarkably, we find that loss of *flr-4* and *flr-1* severely impairs intestinal dipeptide absorption and the intestinal pH gradients that are essential for this process, both of which are restored when TWK-26 is activated. Additionally, we show that the *twk-26* gain-of-function mutation fails to suppress the defects in the defecation motor program displayed by *flr-4* and *flr-1* mutants, demonstrating that nutrient absorption can be uncoupled from the DMP. Our study provides new mechanistic insight into the dual regulation of intestinal metabolism by K^+^ ion homeostasis and protein kinase signaling.

## RESULTS

### A mutation in the TWK-26 potassium channel reverses the metabolic defects conferred by impaired FLR-4 signaling

In *C. elegans*, the intestinal protein kinase FLR-4 promotes growth and maintains lipid homeostasis, partially through regulation of the p38 MAPK pathway (Torzone et al. 2023). Additionally, *flr-4* is required for maximal expression of the intestinal vitellogenin proteins (VIT-1-6), which mediate the transport of lipids to mature oocytes via lipoprotein particles thereby supporting reproduction and embryonic development (Kimble and Sharrock 1983; Torzone et al. 2023). Consistently, loss of *flr-4*, or its kinase activity, results in a striking reduction in body size, growth rate, and intestinal vitellogenin expression.

The downstream targets of FLR-4, which may be other kinases that act in a canonical signaling cascade, are unknown. Expression of the P*vit-3::GFP* vitellogenesis reporter, which consists of the *vit-3* promoter driving GFP (Dowen et al. 2016), is markedly down-regulated after loss of *flr-4*. While adult-specific expression of the vitellogenin proteins is under strict developmental control, their transcriptional regulation is also tied to nutrient sensing mechanisms, as reduced caloric intake decreases the rate of vitellogenin production (Seah et al. 2016; Perez and Lehner 2019). We sought to identify additional components of the FLR-4 pathway and to define whether FLR-4 acts via developmental or nutritional mechanisms to regulate vitellogenesis. To that end, we conducted a forward genetic screen for mutations that suppress the P*vit-3::GFP* expression defects in the *flr-4(ut7)* mutant (Figure 1A). We specifically targeted suppressors in the F1 generation to enrich for gain-of-function suppressor mutations. Following backcrossing to the non-mutagenized strain, we performed whole genome sequencing of the suppressor mutants and identified candidate causative mutations using computational approaches. Remarkably, we identified four, independently generated, instances of the same mutation in *twk-26*, which encodes a two-pore domain potassium (K2P) ion channel (Figure 1A). The *C. elegans* genome encodes 47 K2P channels, most of which have unknown functions (Hobert 2013), including *twk-26*. K2P channels are a class of potassium leak channels that possess four transmembrane domains and two pore-forming domains (Feliciangeli et al. 2015). The TOPCONS protein topology prediction (Tsirigos et al. 2015) for TWK-26 identified four transmembrane domains (Figure 1B), which is consistent with the structure of other K2P channels. The EMS-generated mutation in TWK-26, G485E, is positioned within the C-terminus. The role of TWK-26 in metabolic homeostasis is surprising, as potassium ion leak channels are known to play a role in neurotransmission and vascular tone modulation in mammals, but little is known about potassium channels outside of these excitable tissues (Feliciangeli et al. 2015).

**Figure 1.**
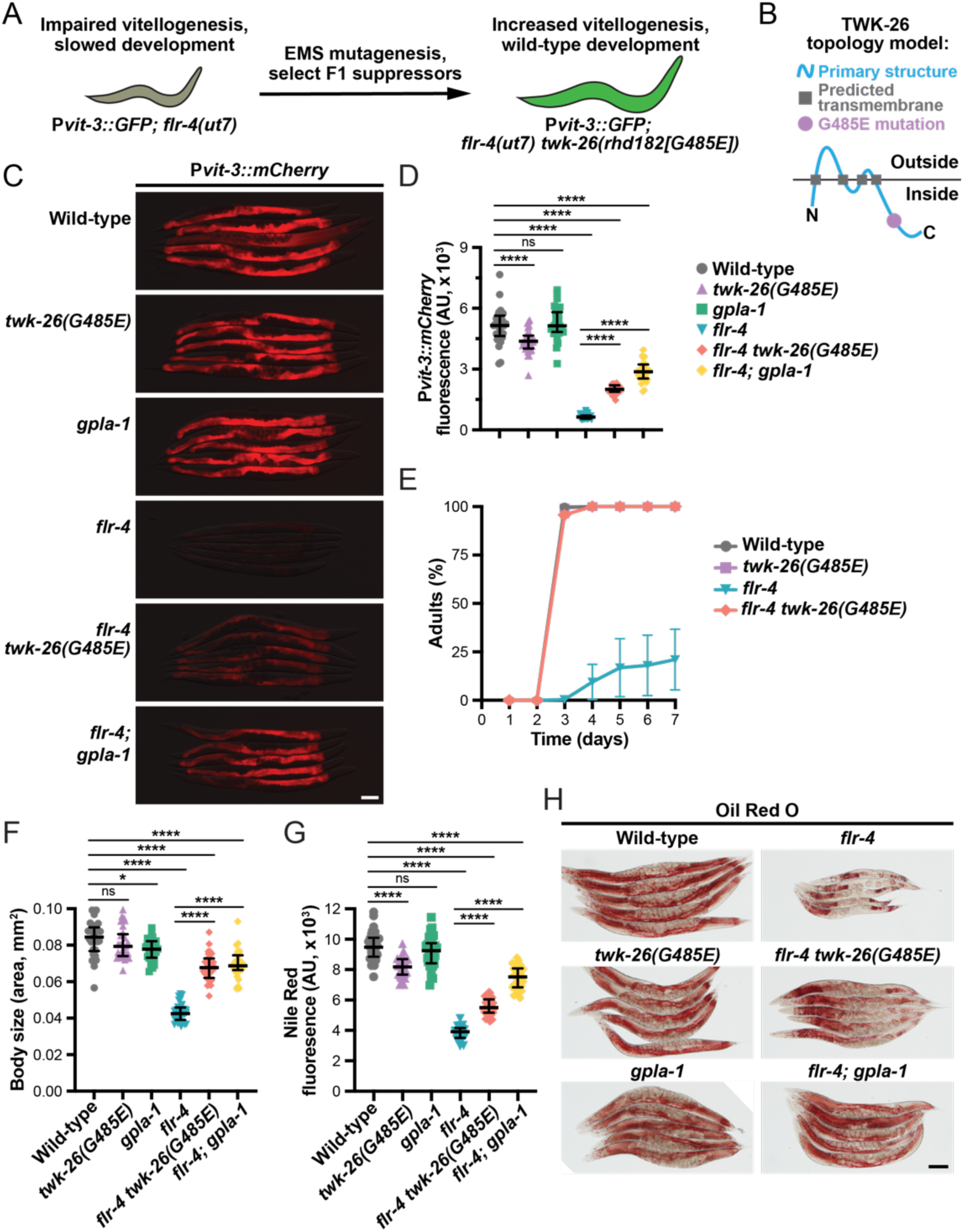
A *twk-26* mutation suppresses the growth and lipid metabolism defects exhibited by the *flr-4* mutant. (A) A diagram illustrating the design of an EMS mutagenesis screen using the *mgIs70[*P*vit-3::GFP]* vitellogenesis reporter. Mutations that suppressed the slow growth or the reporter expression defects of the *flr-4(ut7)* mutant were selected in the F1 generation, thereby enriching for gain-of-function mutations. (B) A transmembrane topology prediction of TWK-26 based on the amino acid sequence. The *twk-26(G485E)* mutation resides in the C-terminal tail. (C) Representative overlaid DIC and mCherry fluorescence images (scale bar, 100μm) and (D) quantification of P*vit-3::mCherry* expression (median and interquartile range; ns, not significant, ****, *P*<0.0001, one-way ANOVA) for the indicated strains at day 1 of adulthood. (E) Growth rate of the indicated strains (mean +/− SEM of 3 independent experiments). Data shown here are also displayed in Figure S1B. (F) Body size of the indicated strains at day 1 of adulthood (median and interquartile range; ns, not significant, *, *P*<0.05, ****, *P* <0.0001, one-way ANOVA). (G) Quantification of Nile Red staining of day 1 adults (median and interquartile range; ns, not significant, ****, *P*<0.0001, one-way ANOVA). (H) Representative images of the indicated strains stained with Oil Red O (day 1 adults; scale bar, 100μm). (C-H) All strains contain the *rhdSi42[*P*vit-3::mCherry]* reporter transgene.

We previously demonstrated that loss of GPLA-1, a neuronally-expressed glycoprotein hormone also known as FLR-2, its receptor, FSHR-1, or downstream cAMP signaling suppressed the growth and metabolic defects displayed by the *flr-4* mutant (Torzone et al. 2023). Using the highly sensitive single-copy P*vit-3::mCherry* transcriptional reporter (Cully et al. 2024), we confirmed that the *twk-26(G485E)* mutation partially suppressed the vitellogenesis defects conferred by the *flr-4* mutation, but not to the same extent as the *gpla-1* mutation (Figure 1C-D). To further elucidate the role of TWK-26 in growth and metabolic homeostasis, we compared the development of the *flr-4(ut7)* single mutant to the *flr-4(ut7) twk-26(G485E)* double mutant. Interestingly, the *twk-26(G485E)* allele strongly suppressed the developmental delay and body size defects conferred by the *flr-4* mutation to levels similar to that of the *gpla-1* allele (Figure 1E-F). Somatic lipid accumulation, as determined by Oil Red O and Nile Red staining, was also partially suppressed by the *twk-26* mutation (Figures 1G-H). Together, these results reveal that TWK-26 plays an important role in regulating growth and lipid homeostasis in conjunction with FLR-4.

The *flr-4(ut7)* allele, as well as mutations in *flr-1* and *flr-3*/*drl-1*, were isolated in a forward genetic screen for mutations that confer resistance to fluoride (Katsura et al. 1994). FLR-1, an acid-sensing sodium ion channel, and FLR-4 are known to colocalize and function together in the intestine (Kobayashi et al. 2011). Furthermore, DRL-1 and FLR-4 form a protein kinase complex in the intestine (Torzone et al. 2023). Thus, we reasoned that if *twk-26(G485E)* suppressed the *flr-4(ut7)* mutation, it may also suppress the *flr-1* and *drl-1* mutations. Indeed, the *twk-26* allele strongly suppressed the growth rate defects of the *flr-1* and *drl-1* mutants, as well as provided a modest suppression of the P*vit-3::mCherry* expression defects (Figure S1A-B). Additionally, the *twk-26(G485E)* mutation suppressed the lipid storage and body size defects of the *flr-1* mutant to similar levels as the *gpla-1* mutation (Figure S1C-D), suggesting that the metabolic and growth defects associated with loss of FLR-1 is due to impaired ion homeostasis. Together, our data indicate that the TWK-26 potassium channel functions in concert with the FLR-4 signaling pathway to coordinate growth and lipid homeostasis in *C. elegans*.

### The *twk-26(G485E)* mutation is a gain-of-function allele

We isolated the *twk-26(G485E)* allele in the F1 generation of an EMS mutagenesis screen, suggesting that the G485E mutation confers either a gain-in-function or haploinsufficiency. To differentiate between these two possibilities, we knocked-down *twk-26* using RNAi in the *flr-4(ut7)* and *flr-4(ut7) twk-26(G485E)* mutants and assessed the resulting phenotypes. If *twk-26(G485E)* is haploinsufficient, *twk-26* RNAi would suppress the growth rate, body size, and metabolic defects of the *flr-4(ut7)* mutant; however, if *twk-26(G485E)* is a gain-of-function allele, *twk-26* RNAi would reverse the suppressive effects that we observed in *flr-4(ut7) twk-26(G485E)* animals. Knock-down of *twk-26* abrogated the suppressive effects of the *twk-26(G485E)* allele, producing animals with reduced developmental rates and small body sizes that are characteristic of the *flr-4* mutant (Figure 2A-C).

**Figure 2.**
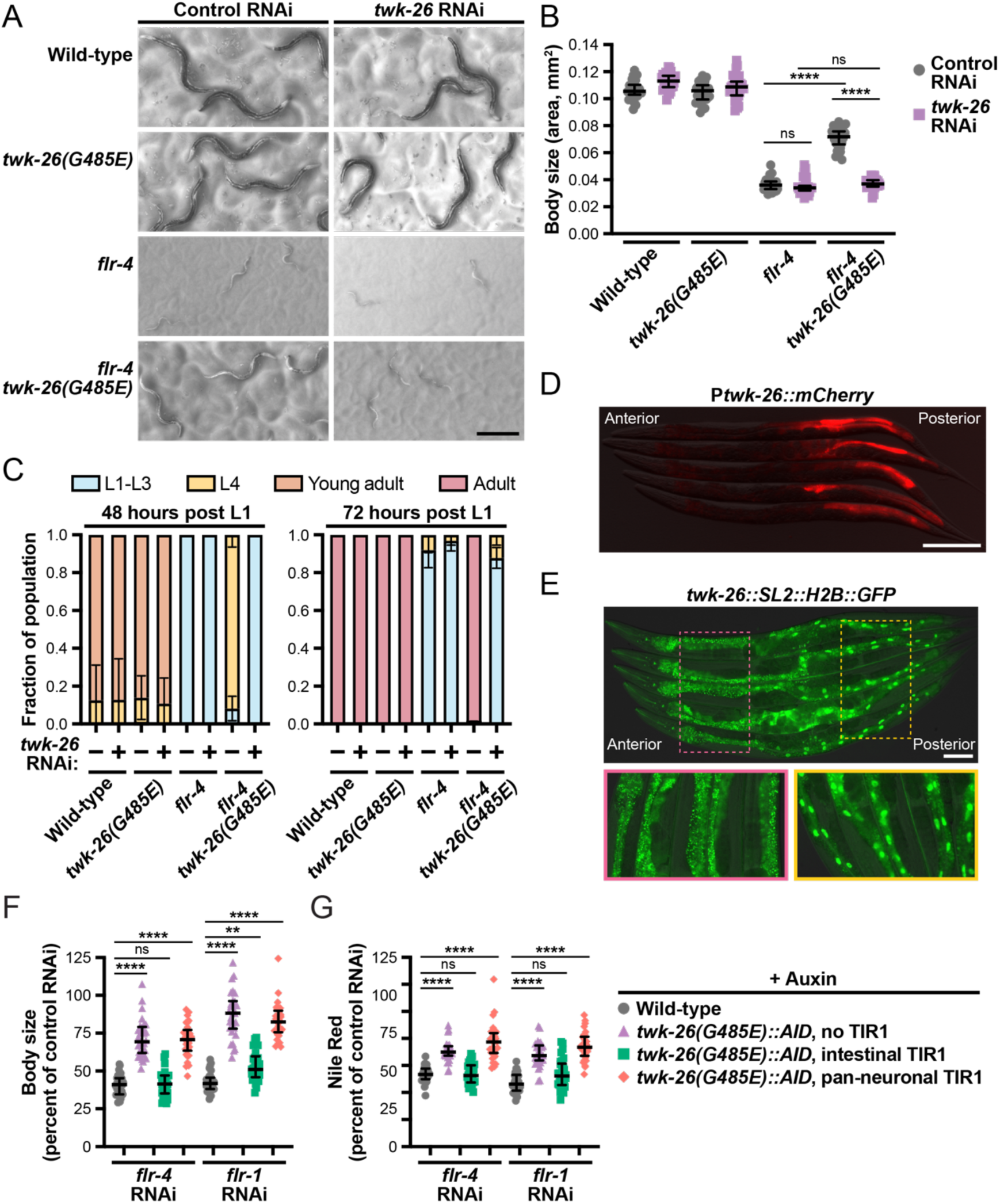
Activation of intestinal TWK-26 suppresses *flr-4* mutant defects. (A) Representative bright field images of the indicated strains subjected to control or *twk-26* RNAi for 72 hours (scale bar, 250μm). (B) Body size of the indicated strains (day 1 adults; median and interquartile range; ns, not significant, ****, *P*<0.0001, one-way ANOVA). (C) Fraction of the population at each developmental stage (mean +/- SD) after treatment of the indicated strains with control or *twk-26* RNAi for 48 or 72 hours. Representative overlaid DIC and fluorescence images of day 1 adult animals carrying (D) a high-copy P*twk-26::mCherry* transgene (scale bar, 200μm) or (E) a *twk-26::SL2::H2B::GFP* allele generated by CRISPR/Cas9 editing (scale bar, 100μm). Quantification of (F) body size or (G) Nile Red staining of the indicated strains subjected to *flr-4* or *flr-1* RNAi following intestinal (P*ges-1::TIR1*) or pan-neuronal (P*rgef-1::TIR1*) depletion of TWK-26::AID protein (4mM auxin) in day 1 adults. All data are normalized to empty vector RNAi controls and displayed as median and interquartile range (ns, not significant, **, *P*=0.0027, ****, *P*<0.0001, one-way ANOVA). (A-C, E, F) All strains contain the P*vit-3::mCherry* reporter.

Although *twk-26* RNAi yielded only a modest effect on P*vit-3::mCherry* expression in *flr-4(ut7) twk-26(G485E)* animals, the *twk-26(G485E)* mutation alone significantly reduced the expression of the vitellogenin reporter, which was reversed by *twk-26* RNAi (Figure S2A). Next, we generated a presumed loss-of-function frameshift mutation within the *twk-26* locus by CRISPR/Cas9 editing and found that this mutation failed to suppress the vitellogenesis defects and small body size of animals depleted for intestinal FLR-4 (Figures S2B and S2C). Together, these data indicate that the *twk-26(G485E)* mutation is a gain-of-function allele, providing unique molecular insight into our understanding of the TWK-26 K2P channel. Thus, we presume that activation, and not inhibition, of the TWK-26 ion channel reverses the dramatic developmental and metabolic defects of the *flr-4* mutant.

We next sought to identify the tissue where TWK-26 functions. It is mostly likely that TWK-26 functions cell-autonomously in the intestine, where FLR-4, FLR-1, and DRL-1 act to promote growth and maintain lipid homeostasis (Kobayashi et al. 2011; Torzone et al. 2023). However, it is possible that TWK-26 functions in neurons, where K2P channels have well-described roles in neurotransmission, locomotion, and innervation of enteric muscles (Gottschling et al. 2017; Yue et al. 2022; Zhou et al. 2022). To define the tissues that express TWK-26, we first generated a P*twk-26::mCherry* transcriptional reporter, which consists of ∼1 kb of the *twk-26* promoter driving mCherry, and overexpressed the transgene in animals as an extrachromosomal array. Surprisingly, we observed mCherry signal specifically in the posterior intestine (Figure 2D), suggesting that *twk-26* expression is repressed in the anterior intestine. To eliminate the possibility of overexpression and transgene artifacts, we used CRISPR/Cas9 editing to insert a SL2::H2B::GFP cassette into the 3’ end of the *twk-26* locus, which generates a polycistronic pre-mRNA that is trans-spliced to generate both the *twk-26* and *h2b::gfp* transcripts. Consistent with our previous results, we observed H2B::GFP expression specifically in the posterior intestine (Figure 2E); however, *twk-26* may also be expressed in a small set of head neurons. To establish whether *twk-26(G485E)* is required in either of these tissues to exert its suppressive effects, we engineered a C-terminal auxin-inducible degron (AID) into the endogenous *twk-26(G485E)* locus by CRISPR/Cas9 editing and performed pan-neuronal or intestine-specific depletion of TWK-26 using auxin. Depletion of TWK-26 G485E in the intestine, but not in neurons, resulted in a decrease in body size and lipid levels in animals lacking *flr-4* or *flr-1* (Figures 2F-G). Together, these data suggest that activation of TWK-26 in the intestine, but not in neurons, restores proper growth and metabolic function upon loss of FLR-4.

### Destabilized gating of the TWK-26 channel restores metabolic homeostasis following loss of *flr-4* or *flr-1*

The predicted human ortholog of TWK-26 is KCNK3/TASK, a two-pore domain K^+^ leak channel that is broadly expressed, including in the brain and small intestine, and that is strongly inhibited by extracellular acidification (Duprat et al. 1997). KCNK3 contains a structural motif called the X-gate, which regulates the opening and closing of the channel (Rödström et al. 2020). We hypothesized that the G485E gain-of-function mutation might confer increased ion conductance of the TWK-26 channel.

Therefore, we explored the possibility that mutations at positions conserved across K2P family members that are known to increase conductance may also impact TWK-26 activity and suppress the *flr-4* mutant defects. A previous study demonstrated that a leucine to asparagine substitution in the second transmembrane domain of several *C. elegans* K2Ps increases channel activity and open probability (Ben Soussia et al. 2019), suggesting that this residue is critical for gating of all K2P channels. Therefore, we engineered the analogous mutation, L229N, into the *twk-26* locus using CRISPR/Cas9 genome editing (Figure 3A). We predicted that if the G485E mutation increases ion conductance of the TWK-26 channel to suppress the metabolic defects of the *flr-4* mutant, then the L229N mutation should confer the same suppressor phenotype. Indeed, the *twk-26(L229N)* mutation suppressed the small body size and lipid storage defects of *flr-4* or *flr-1* knock-down animals to a level similar to that of the *twk-26(G485E)* mutation (Figures 3B-C). This result suggests that the *twk-26(G485E)* mutation identified in our EMS screen confers a gain-in-function that increases conductance through the ion channel. Notably, the L229N mutation was not recovered in our mutagenesis screen; however, the nucleotide conversions preferentially generated by EMS mutagenesis would not be expected to generate this specific amino acid substitution.

**Figure 3.**
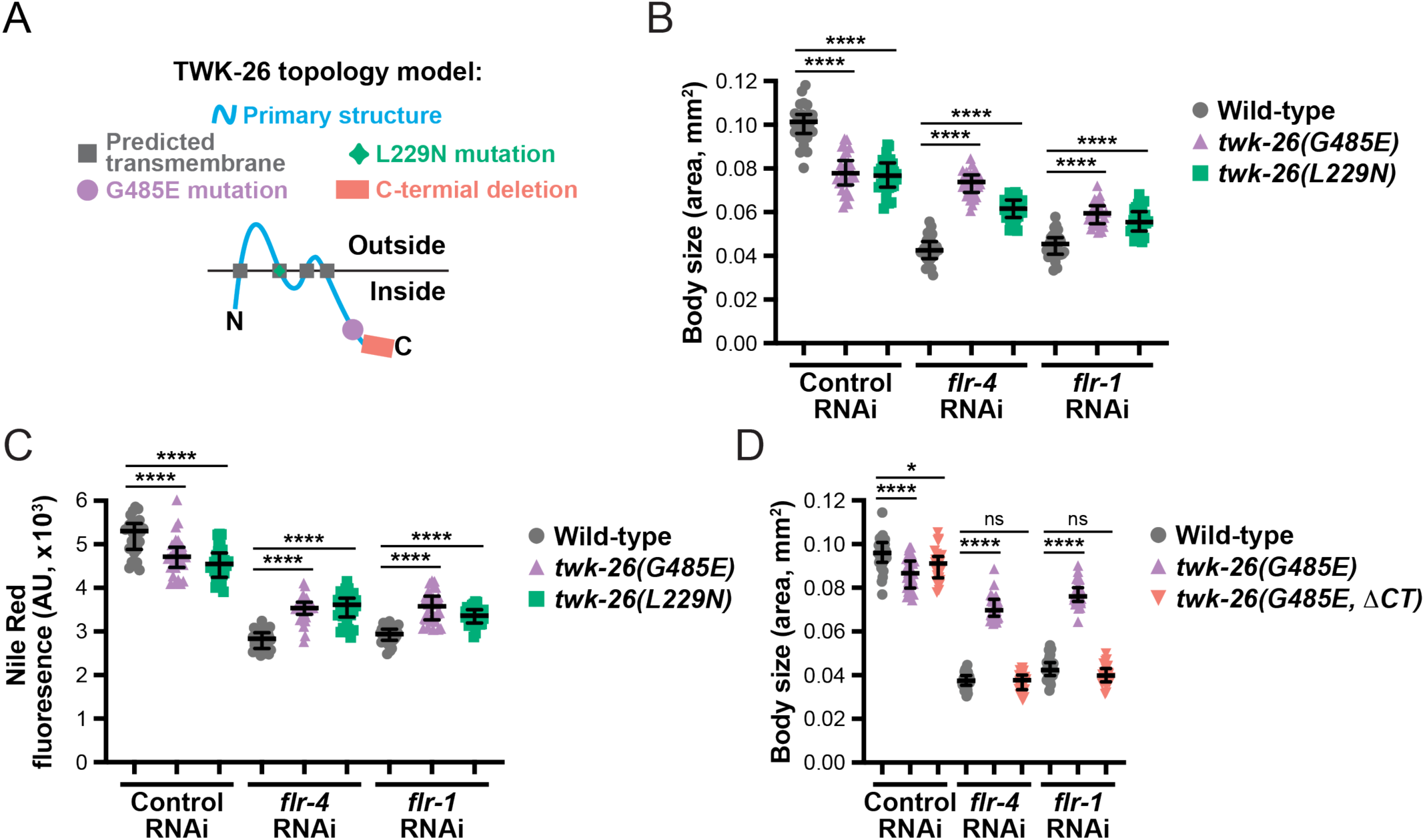
Increased TWK-26 channel gating suppresses the *flr-4* mutation. (A) A transmembrane topology prediction of TWK-26 based on amino acid sequence. The *twk-26(G485E)* mutation resides in the C-terminal tail while the *twk-26(L229N)* mutation is positioned in the second transmembrane domain. The *twk-26(G485E, ΔCT)* strain contains a frameshift at Thr502, resulting in loss of amino acids 503-519. Quantification of (B) body size and (C) Nile Red staining for the indicated strains subjected to control, *flr-4*, or *flr-1* RNAi (day 1 of adults; median with interquartile range; ****, *P*<0.0001, one-way ANOVA). (D) Body size measurements of day 1 adult wild-type or *twk-26* mutant animals carrying the *rhdSi42[*P*vit-3::mCherry]* transgene following control, *flr-4*, or *flr-1* RNAi (median with interquartile range; ns, not significant, *, *P*=0.0136, ****, *P*<0.0001, one-way ANOVA).

Given that the G485E mutation is positioned in the C-terminus of TWK-26, we explored the possibility that the C-terminus of the protein facilitates the increase in TWK-26 G485E activity. Using CRISPR/Cas9 genome editing, we generated a strain expressing the TWK-26 G485E protein lacking the wild-type C-terminus (*i.e.*, ΔSer503-Glu519, frameshift at Thr502). Indeed, the C-terminus of TWK-26 is required for the gain-in-function, as the *twk-26(G485E, ΔCT)* mutation failed to suppress the small body size of *flr-4* or *flr-1* knock-down animals (Figure 3D). These results argue that the C-terminus promotes the gain-of-function activity of TWK-26 G485E; however, we cannot rule out the possibility that this complex mutation impaired the normal function of TWK-26 altogether.

### TWK-26 G485E suppresses autophagy in dietary restriction-like mutants

During dietary restriction (DR), organisms rewire metabolic pathways to initiate catabolic programs, granting access to stored energy reserves when external nutrients are scarce. For example, rapid depletion of intestinal lipids during the initial stages of starvation is a survival mechanism that is conserved across species (Tan et al. 2011). In *C. elegans*, the transcription factor HLH-30, which is orthologous to mammalian TFEB (Lapierre et al. 2013), up-regulates metabolic genes involved in lipolysis during nutrient deprivation (O’Rourke and Ruvkun 2013). Importantly, HLH-30 couples the breakdown of lipid droplet fats with autophagy (O’Rourke and Ruvkun 2013), which shuttles fats from lipid droplets to lysosomes during lipophagy (Singh et al. 2009).

Reduced FLR-4, DRL-1, or FLR-1 signaling strongly induces a DR-like phenotype that is characterized by loss of somatic lipid stores. Therefore, we hypothesized that the *flr-4* and *flr-1* mutants would display an enhanced autophagic response that would be suppressed by activation of TWK-26. To test this hypothesis, we used a dual-fluorescent mCherry::GFP::LGG-1 reporter to visualize the induction of autophagy in *twk-26(G485E)* mutants. The mCherry::GFP::LGG-1 protein, which is orthologous to Atg8, co-labels autophagosomes with mCherry/GFP and labels autolysosomes with only mCherry due to pH quenching of the GFP (Chang et al. 2017). Because an increase in autophagic activity is accompanied by an upregulation of autophagic vesicles (Chang et al. 2017), we quantified the number and size of mCherry-labelled autolysosomes in intestinal cells to assess autophagic flux. We did not observe an appreciable number of mCherry/GFP co-labeled puncta, consistent with a previous study that employed this strain (Chang et al. 2017), and therefore, we did not quantify the number of autophagosomes.

Importantly, quantification of autolysosomes alone is a sufficient measure of autophagic activity, given that the mCherry autolysosomes are known to co-localize with LysoTracker (Chang et al. 2017). We found that knock-down of *flr-4* or *flr-1* by RNAi resulted in a dramatic increase in the amount and size of intestinal autolysosomes (Figures 4A-B), indicating that autophagic activity was up-regulated in these DR-like mutants. Crucially, this increase in autophagy was suppressed by *twk-26(G485E)*, suggesting that activation of the TWK-26 channel suppresses the *flr*-induced DR-like response.

**Figure 4.**
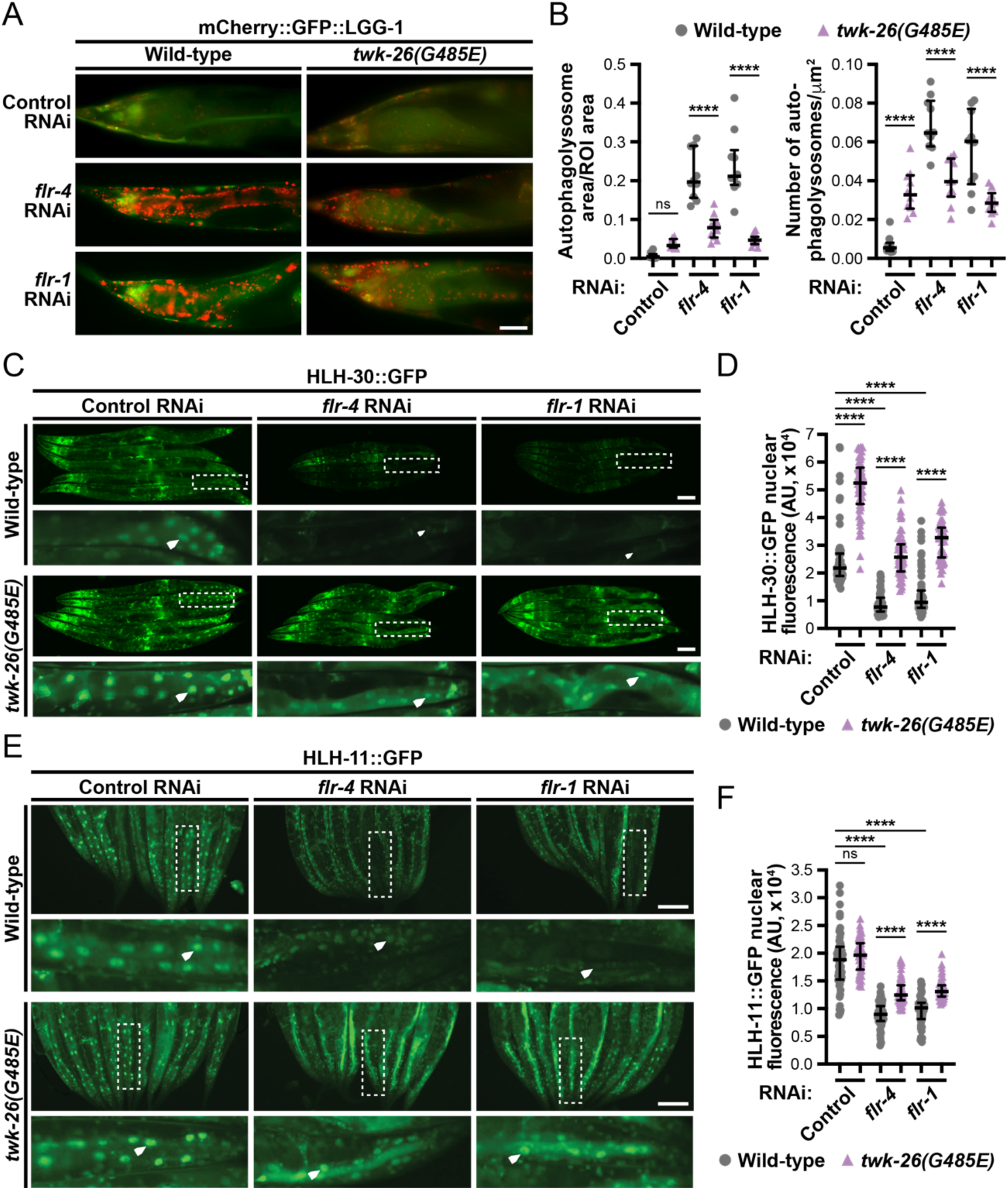
TWK-26 G485E limits autophagy in the *flr-4* and *flr-1* mutants. (A) Representative merged images of mCherry::GFP::LGG-1 puncta in posterior intestinal cells of wild-type or *twk-26(G485E)* day 1 adult animals subjected to control, *flr-4*, or *flr-1* RNAi (scale bar, 25μm). Autolysosomes are labelled by mCherry alone since the GFP signal is quenched. (B) Quantification of the area (left panel) and total number (right panel) of autolysosomes in the posterior intestine (median and interquartile range; ns, not significant, ***, *P*=0.0001, ****, *P*<0.0001, one-way ANOVA). (C) Representative HLH-30::GFP fluorescence images (white arrowheads indicate intestinal nuclei; scale bars, 100μm) and (D) quantification of nuclear HLH-30::GFP signal (median with interquartile range; ****, *P*<0.0001, one-way ANOVA) in wild-type or *twk-26(G485E)* day 1 adults subjected to control, *flr-4*, or *flr-1* RNAi. (E) Representative HLH-11::GFP fluorescence images (white arrowheads, intestinal nuclei; scale bars, 100μm) and (F) quantification of nuclear HLH-11::GFP signal (median with interquartile range; ns, not significant, ****, *P*<0.0001, one-way ANOVA) in wild-type or *twk-26(G485E)* day 1 adult animals following control, *flr-4*, or *flr-1* RNAi.

Given that loss of *flr-4* and *flr-1* increased autophagic flux, we hypothesized that the HLH-30/TFEB transcription factor, which up-regulates autophagy factors during starvation (Settembre et al. 2013), would accumulate in the intestinal nuclei of the *flr* mutants. Surprisingly, knock-down of *flr-4* or *flr-1* by RNAi severely reduced the levels of nuclear HLH-30 (Figures 4C-D), which was suppressed by *twk-26(G485E)*. Consistently, knock-down of *hlh-30* by RNAi had little impact on P*vit-3::mCherry* reporter expression nor the body sizes of the *flr-4* or *flr-4(ut7) twk-26(G485E)* mutants (Figures S3A-B). We also observed that *twk-26(G485E)* alone promoted HLH-30 nuclear localization (Figures 4C-D), suggesting that the animal may activate protective signaling to maximize resources under increased K^+^ channel activity. These results argue that loss of *flr-4* or *flr-1* prompt an HLH-30-independent induction of autophagy or that HLH-30 nuclear localization activates different pathways in different models of dietary restriction.

Therefore, we investigated whether the starvation-responsive transcription factor HLH-11/TFAP4 responds to loss of *flr-4* or *flr-1*. HLH-11 represses the expression of lipid catabolism genes under well-fed conditions and is degraded during starvation to release fat reserves for energy (Li et al. 2020). Indeed, knock-down of *flr-4* or *flr-1* by RNAi reduced levels of nuclear HLH-11 in the intestine; however, this reduction in HLH-11 nuclear signal was suppressed by the *twk-26(G485E)* allele (Figures 4E-F), suggesting that TWK-26 activation prevents the degradation of HLH-11 when nutrients are available.

Together, these data argue that loss of FLR-4 signaling triggers an HLH-11-dependent, and HLH-30-independent, induction of autophagy during dietary restriction. Thus, these data reveal a novel role for the TWK-26 K2P channel in cellular metabolism, as activation of this ion channel partially reverses an autophagic starvation response.

### The TWK-26 G485E mutation uncouples defecation cycle from lipid metabolism

To understand how TWK-26 hyperactivation may be influencing intestinal metabolism to rectify the lipid storage and developmental defects of the *flr* mutants, we first investigated whether the defecation motor program (DMP) is altered in the *twk-26(G485E)* mutant. The *flr-4* and *flr-1* mutants display an irregular defecation cycle (Iwasaki et al. 1995; Katsura et al. 1994; Take-uchi et al. 1998, 2005), resulting in a ‘constipated’ phenotype that may contribute to their DR-like state (Take-uchi et al. 1998). This phenotype, which likely results from improper expulsion of gut contents (Liu and Thomas 1994), coincides with intestinal bloating (Gruss and Corsi 2022; Lin-Moore et al. 2021). Intestine bloating can be indicative of increased microbial colonization, activation of innate immunity pathways, and initiation of pathogen aversion behaviors (Singh and Aballay 2019). Consistently, an impaired DMP is associated with intestinal bloating, enhanced susceptibility to pathogens, and DR-like phenotypes (Das et al. 2023), which could underlie the metabolic and growth defects displayed by the *flr* mutants.

Therefore, we hypothesized that the *flr-4* and *flr-*1 mutants would show intestinal bloating and increased immune signaling resulting from an irregular DMP, which would be suppressed by the *twk-26(G485E)* mutation. We first examined the morphology of the intestinal lumen of animals expressing ERM-1::eGFP, which localizes to the intestinal apical membrane and facilitates lumen width measurements (Ramalho et al. 2020). Consistent with the previously reported DMP defects (Take-uchi et al. 1998, 2005), knock-down of *flr-4* or *flr-1* by RNAi increased intestinal lumen width; however, the bloating was not suppressed by the *twk-26(G485E)* mutation (Figures 5A-B). This result is surprising since intestinal bloating is often associated with a DR-like state (Das et al. 2023), which is displayed by the *flr* mutants and is suppressed by the *twk-26(G485E)* allele (Figures 1G-H and S1D). Furthermore, activation of the p38 innate immune pathway, as assessed by oligomerization of the upstream p38 regulator TIR-1/SARM1 (Peterson et al. 2022), was stimulated by loss of *flr-4* as expected (Torzone et al. 2023); however, TIR-1 aggregation was not suppressed by the *twk-26(G485E)* allele (Figures 5C and S4A-C). We also explored the possibility that the intestinal lumen bloating observed upon loss of *flr-4* or *flr-1* could result from impaired feeding as previously observed in the *phm-2* and *eat-2* mutants, which display food avoidance behaviors and dietary restriction phenotypes that are characteristic of intestinal bloating (Kumar et al. 2019). Using pharyngeal pumping rates to assess feeding behavior (Avery 1993), we found that the *flr-4* and *flr-1* mutants do not display severe feeding defects (Figure S5), as has been observed for the *eat-2* mutant (Raizen et al. 1995; Lakowski and Hekimi 1998). Together, these data reveal that the DR-like state of the *flr-4* and *flr-1* mutants is uncoupled from the intestinal bloating and immunity phenotypes, suggesting that FLR-4 and FLR-1 may influence defecation, and ultimately metabolic homeostasis, more directly.

**Figure 5.**
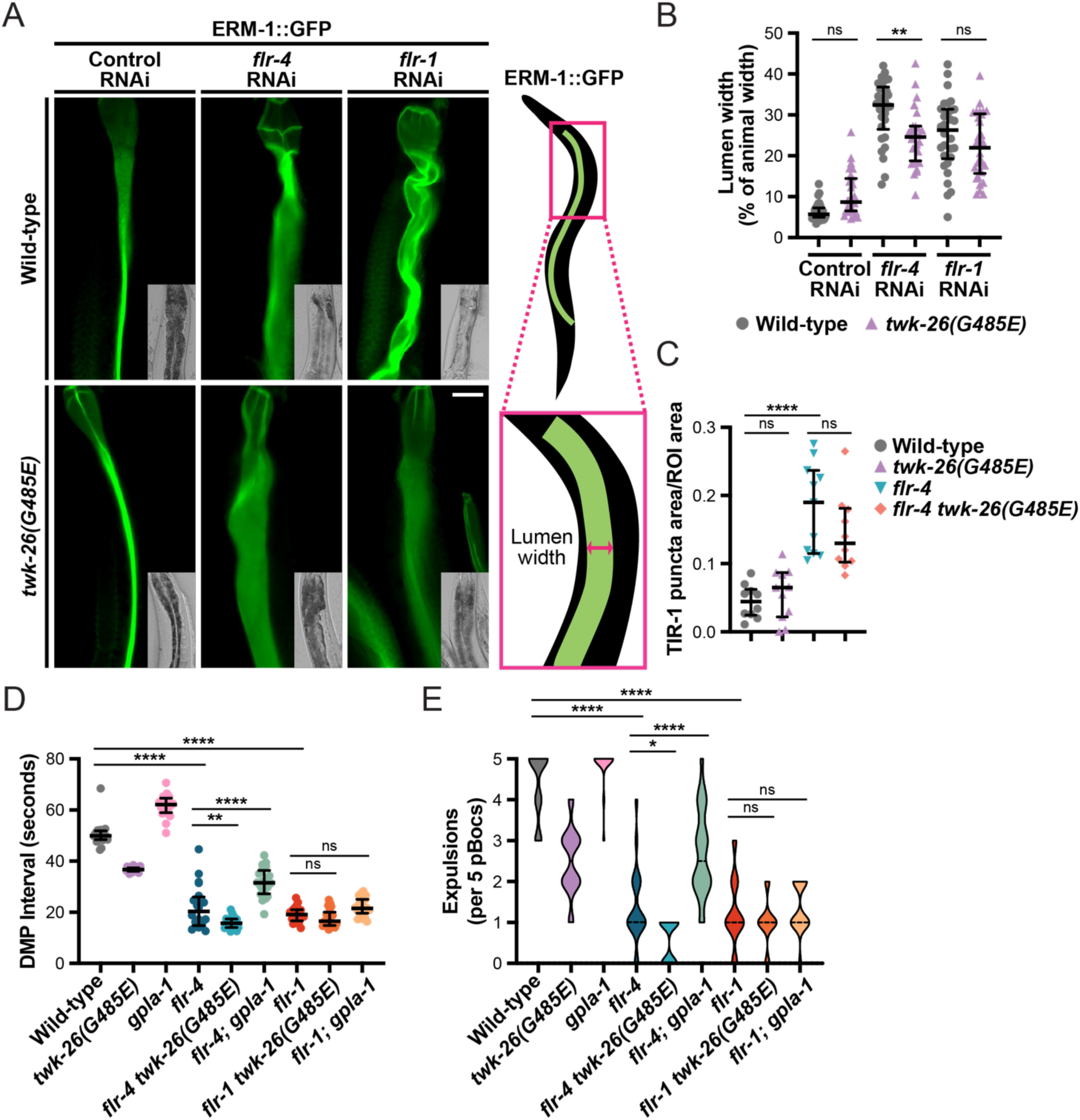
The TWK-26 G485E mutation fails to suppress the intestinal morphology and defecation defects of the *flr-4* and *flr-1* mutants. (A) Representative fluorescence images, as well as DIC insets, of ERM-1::GFP or ERM-1::GFP*; twk-26(G485E)* day 1 adult animals subjected to control, *flr-4*, or *flr-1* RNAi (scale bar, 25μm). The ERM-1::GFP protein marks the intestinal lumen. (B) Quantification of lumen width as a percentage of body width at the measurement site (median and interquartile range; ns, not significant, **, *P*=0.0039, ****, *P*<0.0001, one-way ANOVA). (C) Quantification of TIR-1::wrmScarlet puncta size in the posterior intestine of the indicated strains at day 1 of adulthood (median and interquartile range; ns, not significant, ****, *P*<0.0001, one-way ANOVA). (D) Measurements of the time between posterior body wall contractions (pBoc). Each data point represents the average DMP interval calculated across five pBoc events per individual (median and interquartile range; ns, not significant, **, *P*=0.0014, ****, *P*<0.0001, one-way ANOVA). (E) The number of pBoc events that were accompanied by intestinal expulsion (Exp) across the five observed DMP intervals. Distributions are shown as truncated violin plots with the median value indicated (dashed line). (D-E) Ten animals were scored for two independent experiments for a total of 20 animals per genotype (strains contain *rhdSi42*).

The *C. elegans* DMP is a highly coordinated behavior that is temporally regulated, with a new cycle initiated every 45 seconds and consisting of 3 steps, which together last for 5 seconds (Thomas 1990). These steps include a posterior body wall contraction (pBoc), an anterior body wall contraction (aBoc), and an expulsion, which ultimately results in the release of gut contents. These contractions are reliant on calcium oscillations (Dal Santo et al. 1999; Espelt et al. 2005), which are partially controlled by FLR-1 (Kaulich et al. 2022). Many of the factors that regulate the DMP have been studied in detail; however, it is not clear why *flr-4* and *flr-1* mutations shorten the DMP cycle and cause expulsion defects (Branicky and Hekimi 2006; Iwasaki et al. 1995).

Because defecation is an ion-dependent process, we next explored whether the *twk-26(G485E)* mutation, which likely increases K+ conductance, suppresses the irregularity of the DMP caused by the *flr* mutations by altering intestinal ion flux. The DMP is known to promote absorption of glucose in the intestine (Suzuki et al. 2022), and mutations in key DMP genes, like *flr-1*, restrict fat accumulation (Kaulich et al. 2022). Therefore, it is possible that since the *twk-26(G485E)* allele suppressed the metabolic phenotypes of the *flr-4* and *flr-1* mutants, it may also suppress the irregular defecation rhythm of these animals to indirectly restore fat levels. Surprisingly, we found that *twk-26(G485E)* did not suppress the short DMP interval nor increase the number of intestinal expulsion events in the *flr* mutants (Figures 5D-E). Consistently, knock-down of *nlp-40*, which encodes an intestinal neuropeptide-like protein that controls DVB neuron activity and the expulsion step of the DMP (Wang et al. 2013), resulted in loss of P*vit-3::mCherry* reporter expression and reduced body size; however, these phenotypes were not suppressed by the *twk-26(G485E)* allele (Figure S6A-C). Together, these data indicate that the defects in lipid metabolism are independent of the abnormal DMP in *flr* mutants and further support the notion that TWK-26 has a more direct role in intestinal metabolism.

### TWK-26 G485E restores intestinal dipeptide absorption in the *flr-4* and *flr-1* mutants

Loss of DRL-1 or FLR-4 in the intestine causes a dietary restriction-like (DR-like) state, resulting in small body size, slowed development, and impaired lipid accumulation (Chamoli et al. 2014; Katsura et al. 1994; Torzone et al. 2023). Additionally, loss of *flr-1* results in a DR-like phenotype (Figure S1) (Kaulich et al. 2022; Oishi et al. 2009; Take-uchi et al. 1998). Given the similarity between the mutant phenotypes, it is likely that FLR-1, DRL-1, and FLR-4 act in the same pathway to maintain metabolic homeostasis. Moreover, these DR-like phenotypes were partially suppressed by activating mutations in the intestinal TWK-26 potassium channel (Figure 3), suggesting that impaired ion homeostasis may underlie the DR-like phenotypes in these mutants. Given that the DR-like state of the *flr* mutants is not coupled to the DMP, we explored the possibility that FLR-4 and FLR-1 may govern other ion-driven processes in the intestine to coordinate metabolism, which when disrupted, could be restored by activation of TWK-26. Intriguingly, intestinal nutrient absorption is an ion-dependent process, which could be dysregulated in the *flr-4* and *flr-1* mutants, thereby contributing to the DR-like phenotypes. Mammalian intestinal dipeptide transport via the PepT1 H^+^-symporter is enhanced at low extracellular pH (Fei et al. 1994; Liang et al. 1995). Similarly, two *C. elegans* oligopeptide transporters, PEPT-1 and PEPT-2, display proton gradient-driven electrogenic transport (Fei et al. 1998), with the activity of the intestinal PEPT-1 transporter strongly coupled to intracellular acidification. Importantly, loss of PEPT-1, or the downstream intestinal Na^+^/H^+^ exchanger NHX-2, results in developmental delay and starvation phenotypes (Nehrke 2003). Furthermore, loss of the FLR-1 Na^+^ channel impairs intestinal acidification and results in starvation-like phenotypes (Figure S1 and Figure 4) (Kaulich et al. 2022).

Therefore, we reasoned that the *flr-4* and *flr-1* mutants may display defects in peptide absorption due to impaired intestinal ion gradients. To test this hypothesis, we directly assessed amino acid absorption by feeding animals β-Ala-Lys-AMCA, a fluorescent dipeptide that gets taken up in the gut (Meissner et al. 2004). Interestingly, we observed a marked reduction in dipeptide absorption in the *flr-4* and *flr-1* mutants (Figures 6A-B). These results demonstrate, for the first time, that the FLR-4/FLR-1 signaling axis is required for proper dipeptide absorption and provides a possible mechanism for the starvation-like phenotypes that we observed in the *flr* mutants. Critically, the *twk-26(G485E)* mutation strongly suppressed the dipeptide absorption defects in the *flr-4* and *flr-1* mutants, arguing that the TWK-26 K2P promotes nutrient absorption when activated. Notably, loss of *gpla-1* also suppressed the dipeptide absorption defects in the *flr-4* and *flr-1* mutants (Figures 6A-B), suggesting that neuronal signals may regulate the activity of intestinal ion channels involved in amino acid absorption. We also observed an intriguing spatial preference for TWK-26-dependent dipeptide uptake in the anterior intestine when compared to the posterior intestine (Figure 6B), which is surprising given that *twk-26* expression is significantly higher in the posterior intestine (Figures 2D-E). Together, these results suggest that the FLR-4/FLR-1 pathway governs amino acid absorption by coordinating ion homeostasis alongside of the TWK-26 K+ channel, possibly by directly, or indirectly, altering ion flux in the anterior intestine to maintain intracellular acidification along the length of the intestine.

**Figure 6.**
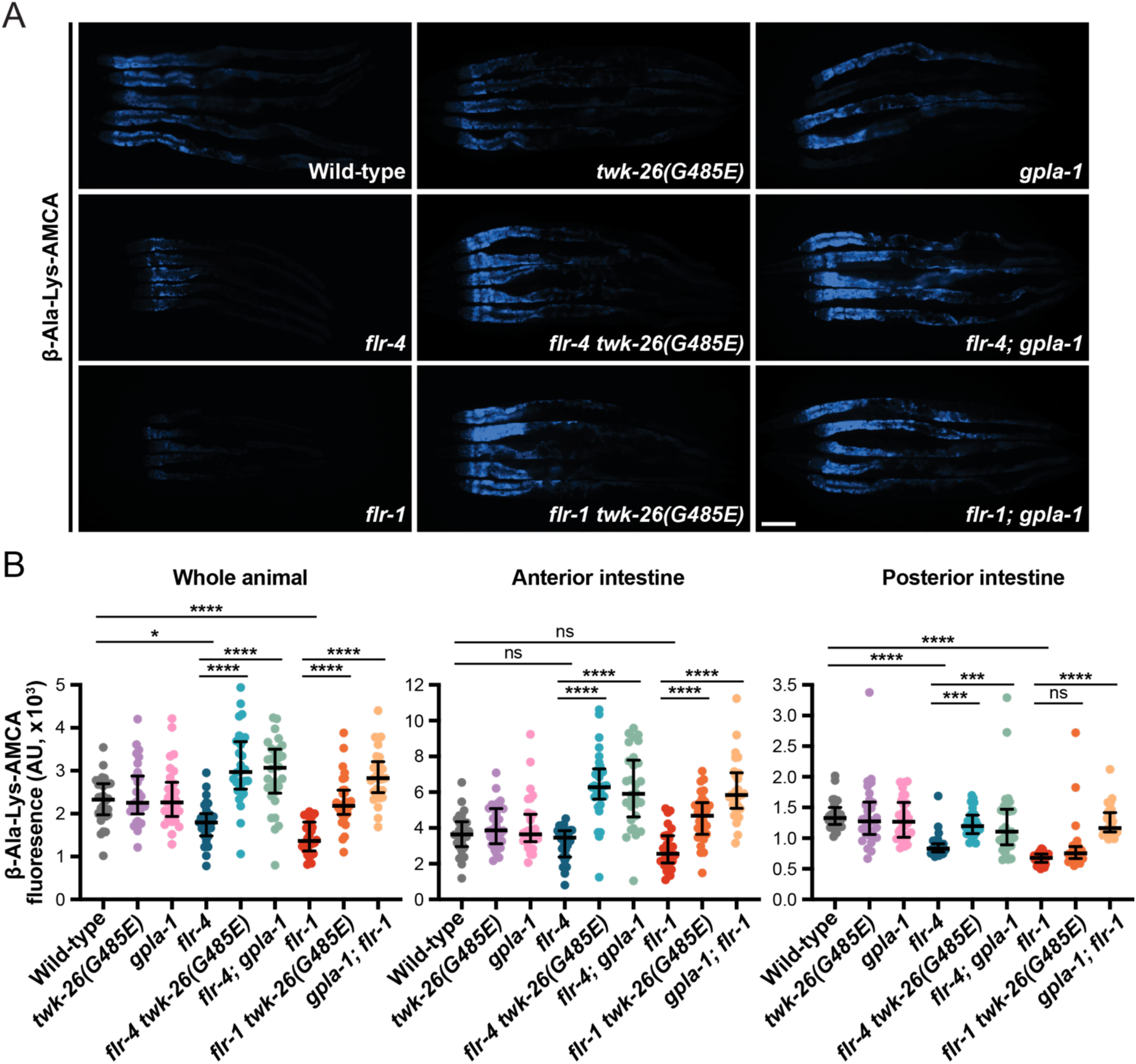
Activation of TWK-26 restores intestinal dipeptide absorption in the absence of *flr-4* and *flr-1*. (A) Representative fluorescence images of day 1 adult animals incubated for 3 hours in 1mM β-Ala-Lys-AMCA, a fluorescently labeled dipeptide (5 animals per image; scale bar, 100μm). Animals are oriented anterior (left) to posterior (right). (B) Quantification of the fluorescent signal in the whole animal or in specified regions of the intestine (anterior region, head to the vulva; posterior region, vulva to the tail). Data are presented as median and interquartile range (ns, not significant, *, *P*=0.012, **, *P*<0.01, ***, *P*<0.001, ****, *P*<0.0001, one-way ANOVA performed for the indicated comparisons).

We could not neglect the possibility that absorption of other nutrients, in particular lipids, may also be dependent on FLR-4, FLR-1, or TWK-26. Surprisingly, a *pept-1* mutation increases lipid uptake, although this may be a compensatory mechanism in response to decreased amino acid absorption (Spanier et al. 2009). To test this hypothesis, we fed animals C1-BODIPY-C12, a fluorescently-labeled fatty acid derivative, to track lipid uptake in the intestine of the *flr-4* and *flr-1* mutants (O’Rourke et al. 2009; Sheng et al. 2015; Spanier et al. 2009); however, we could not clearly resolve a reproducible phenotype and were unable to establish a role for the FLR-4/FLR-1 pathway in lipid absorption using this approach.

Alternatively, we fed Nile Red, a lysochrome dye that accumulates in intestinal lipid droplets and lysosome-related organelles following absorption (O’Rourke et al. 2009), to *flr-4* mutants. We found that *flr-4* mutants displayed significantly less Nile Red signal in the intestine compared to the wild-type control or the *flr-4 twk-26(G485E)* double mutant (Figure S7), indicating that *flr-4* may be required for proper uptake of lipids or lipophilic molecules in the intestine.

### FLR-4 and TWK-26 balance intracellular and luminal pH to regulate nutrient absorption

Loss of either PEPT-1, an intestinal oligopeptide transporter, or NHX-2, the intestinal Na^+^/H^+^ exchanger, impairs amino acid absorption, reduces growth rate, and restricts lipid accumulation (Meissner et al. 2004; Nehrke 2003). Given that activation of TWK-26 suppressed these same defects in the *flr* mutants, we hypothesized that the *twk-26(G485E)* mutation may also suppress loss of *pept-1* or *nhx-2*.

Although knock-down of *pept-1* or *nhx-2* by RNAi resulted in a dramatic reduction in P*vit-3::mCherry* expression and small body sizes, these phenotypes were not suppressed by either the *twk-26(G485E)* or *gpla-1* mutations (Figure S6A-C), suggesting that *twk-26*/*gpla-1* may function genetically upstream of *pept-1*/*nhx-2* or in a different pathway altogether to suppress the metabolic defects of the *flr-4* and *flr-1* mutants. We also tested whether the *flr* pathway was required for *pept-1*/*nhx-2* gene expression using quantitative PCR, which surprisingly revealed that *pept-1* and *nhx-2* expression is strongly upregulated in the *drl-1* or *flr-4* mutants (Figure S6D), possibly via compensatory mechanisms. Together, these results suggest that the FLR-4 pathway may function independently of the canonical regulators of oligopeptide absorption in the intestine.

It has been previously shown that *flr-1* is essential for intracellular intestinal pH oscillations and that dysregulation of proton signaling upon loss of *flr-1* results in reduced fat storage (Kaulich et al. 2022). Thus, we hypothesized that *flr-4*, which functions in the same genetic pathway as *flr-1*, is also required for proper intestinal pH oscillations, and consequently, proton gradient-driven dipeptide absorption and intestinal lipid accumulation. Using the pHluorin sensor to measure intracellular pH (pHi) in the intestine (Nehrke 2003), we found that the *flr-4* mutants displayed normal resting pHi between contractions but relatively weak acidification concomitant with the DMP (Figure 7A-B). This observation contrasts with the relatively acidic pHi observed in *nhx-2* and *vha-6* mutants, which also display DR-like phenotypes (Nehrke 2003; Pfeiffer et al. 2008; Allman et al. 2009). We next wondered if pH oscillations of the intestinal lumen, which are inversely correlated with changes in pHi, are also impacted by loss of *flr-4*. Importantly, the intestinal lumen, which is thought to be the source of protons that drive intestinal acidification and nutrient transport, is known to alkalinize during defecation (Pfeiffer et al. 2008). Thus, we fed animals dextran-conjugated Oregon Green 488, a pH sensitive dye, to measure the fluid pH of the intestinal lumen. We found that the *flr-4* mutant displayed a less acidic luminal pH and blunted luminal pH oscillations in the posterior intestine relative to the wild-type control (Figure S8), consistent with reduced proton-coupled nutrient uptake. In general, when the pBoc contraction is robust, the luminal contents move from the posterior to anterior intestine and the luminal pH is consistent throughout the length of the intestine. Interestingly, in the *flr-4* mutant, the luminal pH oscillations were restricted to the posterior segment, with little movement of luminal contents, resulting in a pH gradient during contraction (Figure 7C).

**Figure 7.**
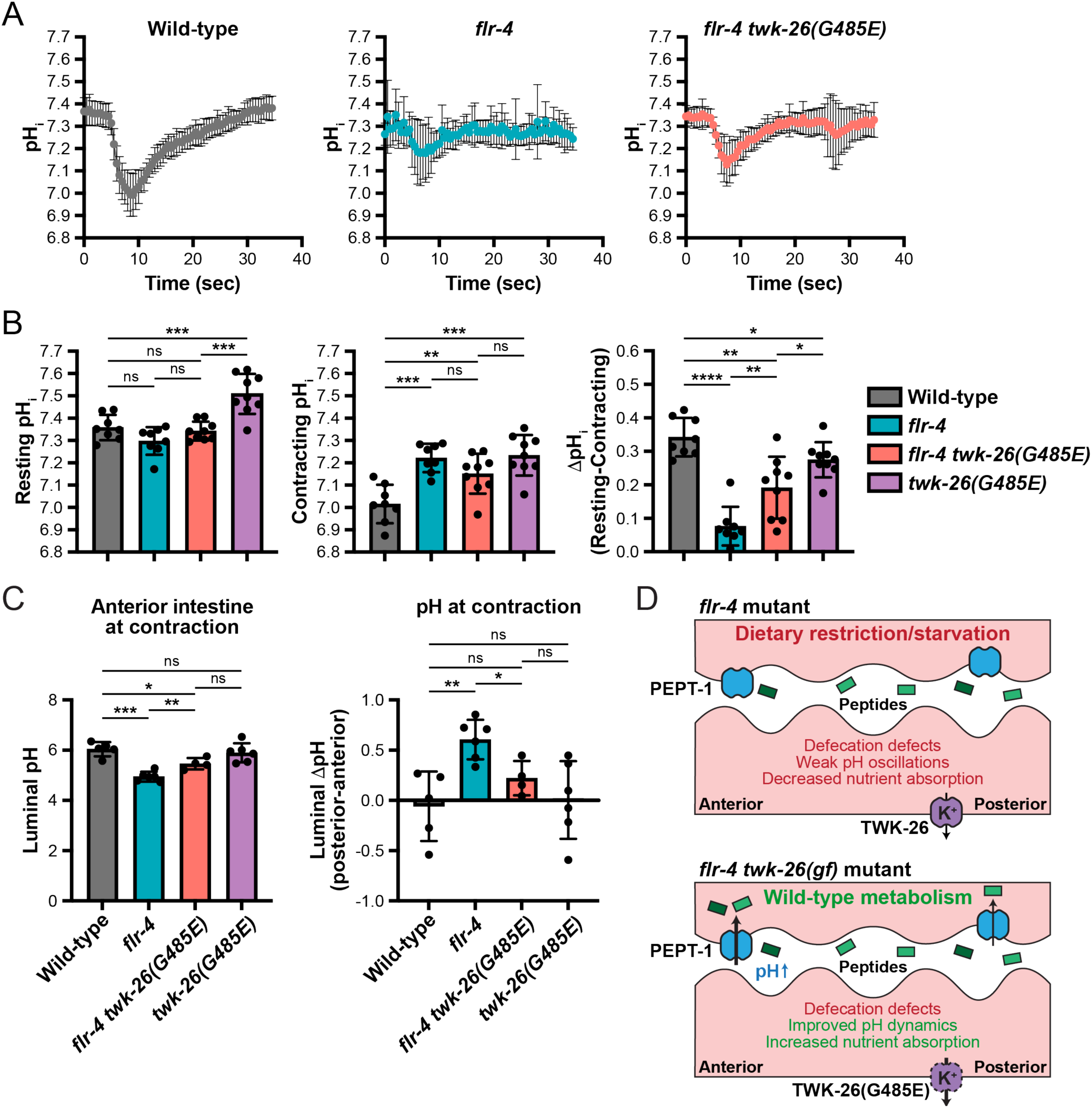
The *flr-4* mutant displays defects in intestinal pH oscillations which are rectified by TWK-26 G485E. (A) A time course (relative to pBoc) of the intracellular pH (pH_i_) of wild-type (N=8), *flr-4(ut7)* (N=9), and *flr-4(ut7) twk-26(G485E)* (N=9) day 1 adult animals expressing the pHluorin transgene. Data are plotted as the mean and SD. (B) Mean resting pH_i_ (left panel) and contracting pH_i_ (middle panel) calculated using the Boltzmann calibration, as well as the ΔpH_i_ (right panel), for each of the indicated pHluorin strains (N=8-9 animals per genotype; mean +/- SD; ns, not significant, *, *P*<0.05, **, *P*<0.01, ***, *P*<0.001, ****, *P*<0.0001, Brown-Forsythe and Welch ANOVA followed by Fishers LSD Test). (C) Measurements of the anterior luminal pH at the time of intestinal contraction in animals fed dextran-conjugated Oregon Green 488 (left panel). The difference in luminal pH between the posterior and anterior intestine is also shown (right panel). All data are presented as mean and SD (N=4-7 animals per genotype; ns, not significant, *, *P*<0.05, **, *P*<0.01, ***, *P*<0.001, Brown-Forsythe and Welch ANOVA followed by Fishers LSD Test). (D) A model depicting how TWK-26 G485E may restore nutrient absorption and metabolic homeostasis upon loss of *flr-4*.

We then asked if activation of TWK-26 can resolve the defects in pH oscillations that we observed in the *flr-4* mutants. While *twk-26(G485E)* was unable to suppress either resting or contracting pHi defects of the *flr-4* mutant when measured independently, it was able to partly rescue the amplitude of the pHi oscillations (Figure 7A-B). More notably, *twk-26(G485E)* suppressed some of the intestinal lumen pH abnormalities in *flr-4* mutants. Specifically, activation of TWK-26 promotes alkalinization of the lumen in the anterior (Figure 7C) but not in the posterior intestine (Figure S8). This decreases the difference between the pH of the anterior and posterior lumens during defecation (Figure 7C), restoring a more wild-type phenotype in the *flr-4* mutant. Together, these data reveal that both *flr-4* and *twk-26* are crucial for regulating pH dynamics of intestinal cells and the gut lumen, providing new insight into the mechanisms by which animals absorb nutrients and maintain metabolic homeostasis.

## DISCUSSION

In this study, we establish a role for the TWK-26 potassium channel in governing lipid homeostasis by promoting intestinal oligopeptide absorption in concert with the FLR-4 signaling pathway. Loss of the FLR-4 kinase, or the downstream FLR-1 sodium channel, results in slow growth, small body size, and reduced lipid accumulation, which likely results from impaired amino acid absorption. This nutrient absorption defect causes a starvation-like state where the HLH-11/TFAP4 transcription factor is likely degraded to initiate catabolic programs in the intestine. Remarkably, the DR-like state of the *flr-4* and *flr-1* mutants is almost completely reversed by an activating mutation in the TWK-26 potassium leak channel. Moreover, our study reveals a new mechanism by which the FLR-4 pathway and the TWK-26 potassium channel together coordinate oligopeptide absorption by altering intracellular and luminal pH dynamics to maintain lipid homeostasis and license proper development.

In genetic models of dietary restriction, it is crucial to understand the specific molecular mechanisms that reduce caloric intake and alter organismal physiology. Mutations in the DRL-1/FLR-4 signaling pathway confer DR-like phenotypes; however, the cause of these phenotypes remains poorly understood (Chamoli et al. 2014; Take-uchi et al. 2005). The metabolic reprogramming stemming from loss of *drl-1* includes increased beta-oxidation, reduced fat storage, and increased lifespan, which are all characteristics of dietary restriction; however, the underlying mechanism has remained elusive (Chamoli et al., 2014). Similarly, loss of FLR-4, which binds to DRL-1 in a complex (Torzone et al., 2023), stimulates cytoprotective and detoxification pathways, which has also been observed in *drl-1* knock-down animals (Chamoli et al. 2014, 2020; Verma et al. 2018).

Consistently, we demonstrate that loss of *flr-4* or *flr-1* results in a dramatic induction of autophagy in the intestine, which has been previously observed during nutrient deprivation (Chamoli et al. 2014; Hansen et al. 2008). HLH-30/TFEB is a conserved transcriptional regulator of autophagy that promotes lysosomal biogenesis (Lapierre et al. 2013; Settembre et al. 2011). In the presence amino acids, mTORC1 is activated and phosphorylates TFEB, preventing its nuclear localization and restricting the induction of autophagy programs (Martina et al. 2012; Roczniak-Ferguson et al. 2012; Sparta et al. 2023). It is possible that signaling through mTORC1 is diminished in *flr-4* and *flr-1* mutants due to impaired absorption of dipeptides (Hara et al. 1998). However, upon loss of *flr-4* and *flr-1*, we observed a decrease in nuclear accumulation of HLH-30, which suggests that either mTORC1 signaling remains active or that *hlh-30*/TFEB is transcriptionally or post-translationally down-regulated in these mutants through a different mechanism. We find that HLH-11 levels are also severely decreased in the *flr* mutants. HLH-11 represses the expression of lipid catabolism genes under fed conditions, causing lipid accumulation; however, HLH-11 is degraded by proteosome upon starvation, promoting lipid utilization (Li et al. 2020). An intriguing possibility is that both HLH-30 and HLH-11 are degraded in response to impaired FLR-4 signaling, resulting in a nuanced transcriptional response that promotes catabolism and up-regulation of lysosomal function; however, it is unclear which transcription factor(s) might contribute to this response. Still, it remains critical to understand the mechanisms underlying this starvation-like response triggered by loss of FLR-4 signaling.

Our discovery that activation of the TWK-26 potassium channel reversed the metabolic defects in the *flr-4* and *flr-1* mutants provided, for the first time, an avenue for exploring the underlying mechanism of the DR phenotypes associated with these mutations, as well as revealed how FLR-4 signaling maintains metabolic homeostasis by governing ion-mediated processes in the intestine to promote growth. Furthermore, we established a novel intestinal function for the TWK-26 leak channel, which has not been previously characterized. KCNK3 is the mammalian ortholog of TWK-26 based on sequence similarity. In mammals, KCNK3/TASK-1 is expressed in neurons (Talley et al. 2001) and forms homodimers or heterodimers with other acid-sensitive TASK channels (Berg et al. 2004), such that these dimeric channels are activated by extracellular alkalization and inhibited by acidification. Inhibition of TASK channels like KCNK3, or its heterodimeric partner KCNK9, results in excitation of sensory neurons (Bautista et al. 2008). These acid-sensitive channels have also been described to regulate chemosensation in the carotid body, which maintains normal respiratory ventilation (Trapp et al. 2008). Thus, TASK channels, like KCNK3, play a crucial role in sensing changes in their environment to maintain homeostasis. While KCNK3 is known to be expressed in the human intestine (Duprat et al. 1997), a functional role in nutrient absorption has not yet been described. Thus, our findings on TWK-26 provide the molecular framework to dissect the metabolic role of KCNK3 in humans.

Furthermore, we elucidated the mechanism by which the TWK-26 potassium channel maintains metabolic homeostasis. We demonstrate that the G485E mutation, which genocopies the gain-of-function TM2.6 mutation (Ben Soussia et al. 2019), likely increases TWK-26 channel activity to promote robust oligopeptide absorption, a function that has not been previously ascribed to KCNK3. It remains elusive how the G485E mutation activates the TWK-26 channel, especially given that the C-terminus is necessary for TWK-26 G485E activity. We have identified residues in the C-terminus that may serve as potential phosphorylation sites (O’Kelly et al. 2002), so it is possible that phosphorylation of the TWK-26 C-terminus is required for channel activation.

It is commonly accepted that the *C. elegans* intestine, which is comprised of 20 epithelial cells, is simplistic and uniform, given that each intestinal cell has arisen from a single progenitor (Zhang et al. 2017; Sulston et al. 1983; Asan et al. 2016). We found that expression of *twk-26* is restricted to the posterior intestine. Little is known about anterior-posterior gene expression patterning in the *C. elegans* intestine, although some studies suggest that this spatial patterning may be governed by Wnt signaling (Schroeder and McGhee 1998; Fukushige et al. 2005). Intriguingly, expression of the *pho-1* acid phosphatase, which is required to promote proper growth, is also limited to the posterior intestine (Fukushige et al. 2005). Thus, it is possible that anterior-posterior gene expression patterning in the intestine governs nutrient absorption differently along the length of the intestine. In support of this idea, activation of TWK-26 strongly promotes dipeptide absorption in the anterior intestine while the *flr-4* and *flr-1* mutants display a decrease in oligopeptide absorption in the posterior intestine. It is possible that the activity of TWK-26 G485E in the posterior intestine is sufficient to promote mixing of luminal contents throughout the length of the intestine, thereby driving robust dipeptide absorption in the anterior intestinal cells.

Consistent with this idea, we found that activation of TWK-26 promotes alkalinization in the lumen of the anterior intestine and partially suppresses the impaired intestinal acidification observed in the *flr-4* mutant. Nutrient absorption in the intestine is driven by a pH gradient across the apical membrane, where acidification of the intestinal lumen enhances the uptake of dipeptides and increases fat accumulation (Allman et al. 2009; Nehrke 2003). Yet, it remains unclear how anterior lumen alkalinization is functionally tied to the robust oligopeptide absorption that we observe upon activation of TWK-26. It is possible that TWK-26 has indirect effects on these intestinal ion gradients. Notably, extracellular alkalinization has been shown to activate other TASK channels (Lotshaw 2006; Reyes et al. 1998; Niemeyer et al. 2007), which may more directly stabilize membrane ion gradients necessary for proper nutrient absorption.

While the ion gradients that regulate the defecation motor program have been previously linked to fatty acid absorption (Spanier et al. 2009), we found that TWK-26 is more directly involved in nutrient absorption and that its activity can be uncoupled from the defecation motor program (DMP). Indeed, activation of TWK-26 failed to suppress the defecation defects of the *flr* mutants, namely the shortened defecation interval and expulsion defects. This result is surprising, given that the DMP plays a central role in nutrient absorption, proper growth, and lipid metabolism (Kaulich et al. 2023; Suzuki et al. 2022). Moreover, TWK-26 activation also failed to suppress the intestinal bloating phenotype of the *flr* mutants, which has been observed in other DR mutants (Das et al. 2023; Katsura et al. 1994; Long et al. 2002). Notably, mTOR pathway mutants also display luminal bloating (Long et al. 2002; Meissner et al. 2004), which is further enhanced by mutation of the dipeptide transporter *pept-1* (Meissner et al. 2004), suggesting that intestinal distension may be a consequence of nutrient deprivation. Intestinal bloating is also observed during pathogen infection (Irazoqui et al. 2010; Singh and Aballay 2019), which upregulates an immune response and alters feeding behaviors in *C. elegans*. Immune signaling through the p38/PMK-1 MAPK pathway is hyperactive in the *drl-1* mutant, restricting growth and down-regulating vitellogenesis (Torzone et al. 2023). Consistently, we observed TIR-1 aggregation in the *flr-4* mutant. However, hyperactivation of TWK-26 does not suppress TIR-1 puncta formation nor intestinal bloating, suggesting that restoration of ion homeostasis in the intestine promotes lipid accumulation and growth independently of altering innate immunity pathways. Excitingly, our findings uniquely uncouple intestinal distension and the DMP from metabolic homeostasis, suggesting that animals may be able to dynamically alter the expression of factors that maintain ion homeostasis to promote efficient uptake and utilization of nutrients.

The FLR-4 pathway supports growth and reproduction by altering intracellular and intestinal lumen pH dynamics to promote the absorption of oligopeptides in the intestine. When signaling through this axis is impaired, animals fail to develop at a normal rate and exhibit a reduction in fat storage and maternal vitellogenin provisioning. Importantly, hyperactivation of the TWK-26 potassium leak channel restores intestinal homeostasis to support normal growth by increasing nutrient absorption, likely by balancing the ion gradients that govern proton-driven dipeptide transport. This work establishes a platform to investigate how protein kinase signaling maintains metabolic homeostasis to promote development and enhances our understanding of how ion channels govern lipid metabolism, which could be important in unraveling the mechanisms of metabolic diseases like diabetes and inflammatory bowel disease.

## METHODS

### C. elegans strains

*C. elegans* strains were maintained on NGM media plates seeded with *E. coli* OP50 at 20**°**C as previously described (Brenner 1974). Animals were synchronized at the L1 larval stage by isolating embryos via bleaching followed by incubation overnight at room temperature in M9 buffer. Auxin inducible degradation (AID) experiments were performed using plates containing 4mM naphthaleneacetic acid (K-NAA, PhytoTech). All strains used in this study are listed in Supplementary Table S1.

### Generation of transgenic animals

To generate the P*twk-26::mCherry::unc-54* transgene, the promoter of *twk-26* (chromosome X: 3,055,393 – 3,054,414; WS295) was amplified from genomic DNA and fused to a *mCherry::unc-54 3’UTR* fragment via PCR fusion (Hobert 2002). The resulting PCR fragment was microinjected *unc-119(ed3)* animals at 20ng/μl, along with 5ng/μl P*myo-3::GFP*, 20ng/μl pCFJ151, and 55ng/μl of 2-Log DNA ladder (New England BioLabs), to generate the DLS1045 strain. Transgenic animals were imaged on a Nikon SMZ-18 Stereo microscope equipped with a DS-Qi2 monochrome camera. Additional independent array lines were generated and displayed a similar distribution of mCherry signal in the intestine.

### CRISPR/Cas9 gene editing

Genomic editing was conducted by germline microinjection of Cas9::crRNA::tracrRNA complexes (Integrated DNA Technologies) as previously described (Ghanta and Mello 2020). crRNA sequences are listed in Supplementary Table S2. Missense mutations were generated using single-stranded oligodeoxynucleotides as the repair donor molecule. To create the *twk-26* C-terminal deletion, a crRNA was designed for each end of the deletion site and a single-stranded repair oligo bridged the repair site, which resulted in a frameshift mutation at Thr502. To insert the 2xmIAA7::3xHA or the SL2::H2B::GFP cassette into the 3’ end of the *twk-26* locus, dsDNA donor molecules with ∼40 bp homology arms were prepared by PCR using Q5 DNA Polymerase (New England BioLabs) and were purified with HighPrep PCR Clean-up beads (MagBio) per the manufacturer’s instructions. Prior to microinjection, the DNA repair template was melted and reannealed (Ghanta and Mello 2020).

### RNAi experiments

*E. coli* HT115(DE3) strains carrying the control pL4440 vector or gene-specific RNAi plasmids (Ahringer RNAi library) were grown for ∼20 hours in Luria-Bertani media with 100μg/mL ampicillin at 37°C. Bacteria were concentrated 30x via centrifugation and seeded onto NGM plates containing 5mM isopropyl-β-D-thiogalactoside (IPTG) and 50μg/ml ampicillin. Plates were maintained at room temperature overnight to induce dsRNA expression. Synchronized L1 animals were dropped on RNAi plates and were grown to day 1 of adulthood.

### Oil Red O staining

Approximately 150 L4 animals were picked to new plates, grown for 24 hours, and harvested as day 1 adults in S-basal media. Animals were washed in S-basal, fixed with 60% isopropanol, and stained for 7 hours with 0.5% Oil Red O as previously described (Dowen 2019). Animals were mounted on 2% agarose pads and imaged using a Nikon SMZ-18 Stereo microscope equipped with a DS-Qi2 monochrome camera for intensity analyses or a Nikon Ti2 widefield microscope equipped with a DS-Fi3 camera for representative color images. Quantification of Oil Red O staining was conducted by manually outlining individual animals in Fiji (Schindelin et al. 2012) and then measuring the mean gray value within the area. These values were subtracted from the maximum gray value for 16-bit images, 65,526, as previously described (DuMez-Kornegay et al. 2024). Data were plotted and analyzed using Prism 9. A one-way ANOVA with a Bonferroni correction was used to calculate *P* values.

### Nile Red staining

A Nile Red stock solution was prepared in acetone at 0.5mg/mL and stored at 4°C. Approximately 150 L4 animals were picked to new plates and harvested 24 hours later as day 1 adults in S-basal media. Animals were washed, fixed in 60% isopropanol, and stained for 2 hours in 0.03mg/mL Nile Red. Animals were mounted on 2% agarose pads and imaged with a Nikon Ti2 widefield microscope equipped with a Hamamatsu ORCA-Fusion BT camera. Quantification of fluorescence intensity was conducted by manually outlining individual animals in Fiji and measuring the average pixel intensity within the area. Values were plotted, and statistical analyses were performed, using Prism 9. A one-way ANOVA was conducted with Bonferroni corrections to calculate *P* values.

### Growth and body size assays

For developmental growth rate assays, animals were grown for several generations without starvation prior to being assayed. Approximately 100-200 freshly laid embryos (20 hours or less) were picked to new plates and scored every 24 hours for 7 days. Gravid adults were removed from the population and were recorded as having reached adulthood. For the single time point developmental assays (Figure 2A and 2C), animals were grown from synchronized L1s on RNAi plates and the developmental stage for each worm was recorded at the indicated time point across the entire population.

To quantify body size, animals were picked at the L4 larval stage to new plates and maintained for 24 hours prior to imaging. Day 1 adult animals were mounted on 2% agarose pads with 25mM levamisole and imaged using a Nikon SMZ-18 stereo microscope equipped with a DS-Qi2 monochrome camera. Body size values were obtained by manually outlining individual worms in Fiji, the area measurements (pixels) were converted to mm^2^ using known imaging parameters, and the data were plotted and analyzed in Prism 9 (one-way ANOVA with a Bonferroni correction for multiple testing).

### Quantitative PCR

Preparation of total RNA and qPCR assays were performed exactly as previously described (Torzone et al. 2023). All primer sequences are listed in Supplementary Table S3.

### EMS Mutagenesis

Mutagenesis of *flr-4(ut7); mgIs70[*P*vit-3::GFP]* animals was performed using 47mM ethyl methanesulfonate (EMS, Sigma-Aldrich) and was conducted as previously described (Dowen et al. 2016). Approximately 330,000 haploid genomes were screened on *flr-4* RNAi plates, which prevented isolation of intragenic suppressor mutations. The *flr-4* suppressor strains were isolated in the F1 generation based on increased growth rate and/or P*vit-3::GFP* reporter expression relative to the non-mutagenized *flr-4(ut7); mgIs70* strain. Suppressor strains were backcrossed to the parental strain four times, the genomic DNA was isolated using the Qiagen Gentra Puregene Tissue Kit (Doitsidou et al. 2010), and libraries for whole genome sequencing were prepared and sequenced by Novogene (Sacramento, CA) using an Illumina NovaSeq instrument (PE 150). Candidate suppressor mutations were identified using in-house scripts, as previously described (Minevich et al. 2012).

### Reporter imaging and quantification

Strains carrying either the high-copy *mgIs70[*P*vit-3::GFP]* transgene or the single-copy *rhdSi42[*P*vit-3::mCherry]* transgene have been previously described (Dowen et al. 2016; Torzone et al. 2023). P*vit-3::mCherry* fluorescence was measured in strains containing the *rhdSi42* reporter. Animals were grown to the L4 stage, picked to new plates, and imaged 24 hours later as day 1 adults on a 2% agarose pads containing 25mM levamisole. A Nikon SMZ-18 Stereo microscope equipped with a DS-Qi2 monochrome camera was used for imaging. In Fiji, the body of each animal was manually outlined in the bright field channel and the mean intensity values were obtained in the mCherry channel. The values were plotted in Prism 9 and a one-way ANOVA with a Bonferroni correction for multiple testing was conducted to obtain *P* values.

To determine the cellular localization of the HLH-30::GFP and HLH-11::GFP transcription factors, strains containing the *sqIs19[HLH-30::GFP]* or *wgIs396[HLH-11::TY1::EGFP::3xFLAG]* transgenes were grown on RNAi plates until the L4 stage, picked to fresh RNAi plates, and imaged 24 hours later as day 1 adults on 2% agarose pads with levamisole using a Nikon Ti2 widefield microscope equipped with a Hamamatsu ORCA-Fusion BT camera. A 10x objective was used to capture representative images and a 20x objective was used to image animals for fluorescence measurements. For each genotype, quantification of nuclear fluorescence was performed by outlining the 2 posterior-most intestinal nuclei for 30 animals and measuring the mean intensity value in Fiji. Mean fluorescence intensity values were plotted in Prism 9 and a one-way ANOVA with a Bonferroni correction for multiple testing was conducted to obtain *P* values.

The *sqIs11[mCherry::GFP::LGG-1]* reporter was used to measure autophagic lysosomes (Chang et al. 2017). Imaging was conducted using a CFI Apo 60X Oil TIRF objective on a Nikon Ti2 widefield microscope equipped with a Hamamatsu ORCA-Fusion BT camera. Animals were grown to the L4 stage on RNAi plates and transferred to freshly made RNAi plates 24 hours prior to imaging. Day 1 adult animals were mounted on 2% agarose pads with levamisole and the posterior-most intestinal cells were imaged in the bright field, mCherry, and FITC channels. Autophagic lysosomes were measured in the posterior intestine where the last 2-4 cells are unobstructed by other tissues. Z-stacks were cropped to 0.6mm to capture planes of the intestine that were easily visualized. Z-stacks were then denoised, deconvoluted, and compressed into a single image using the Nikon NIS-Elements analysis software. Hand-drawn ROIs were placed around the 2-4 posterior-most intestinal cells and mCherry puncta were measured using the object counts feature in NIS-elements. Counts of LGG-1::mCherry puncta and the puncta area relative to the ROI area were plotted in Prism 9 and a one-way ANOVA with a Bonferroni correction was performed to calculate *P* values.

To measure intestinal lumen widths, the *erm-1(mib15[erm-1::eGFP])* strain was used to mark the intestinal lumen (Ramalho et al. 2020). Animals were grown on RNAi plates, picked at the L4 larval stage to fresh RNAi plates, and imaged 24 hours later as day 1 adults mounted on 2% agarose pads containing levamisole. A Nikon Ti2 widefield microscope equipped with a Hamamatsu ORCA-Fusion BT camera was used to image animals at 20x. Intestinal lumen widths were measured in the middle of the animal where the intestine was not obstructed by other tissues. Lumen widths from 30 animals were obtained for each condition, the data were plotted in Prism 9, and a one-way ANOVA with a Bonferroni correction was performed to calculate *P* values.

### Nutrient absorption assays

Nile Red feeding was performed as previously described (Mak et al. 2006) with only minor modifications. Briefly, embryos or L1 animals were picked to Nile Red-containing plates (500μL of 1μg/mL Nile Red solution overlaid onto the bacterial lawns), grown to day 1 of adulthood, mounted on 2% agarose pads containing levamisole, and imaged at 8x zoom using a Nikon SMZ-18 Stereo microscope with a DS-Qi2 monochrome camera using the mCherry and DIC channels.

For dipeptide absorption assays, feeding with β-Ala-Lys-AMCA (BioSynth) was performed as previously described (Meissner et al. 2004). Briefly, animals were grown to the L4 stage, picked to fresh plates, grown for 24 hours, and transferred to 0.5mL Eppendorf tubes containing 1mM β-Ala-Lys-AMCA in 100μL of M9 media. The samples were covered in aluminum foil, incubated for 3 hours at room temperature with gentle rocking, washed three times with M9, and mounted on 2% agarose pads containing levamisole for imaging with a Nikon Ti2 widefield microscope equipped with a Hamamatsu ORCA-Fusion BT camera. Using Fiji, whole animals or a specific body regions (*i.e.*, anterior, posterior) were outlined in the brightfield channel and the mean fluorescence measurements were measured in the DAPI channel. The values were plotted in Prism 9 and a one-way ANOVA with a Bonferroni correction for multiple testing was conducted to obtain the *P* values.

### Defecation assays

Videos of day 1 adult animals were recorded using a Nikon SMZ-18 Stereo microscope with a DS-Qi2 monochrome camera. Animals were permitted to freely move and feed on *E. coli* OP50 NGM plates while recording. One animal at a time was recorded for a total of 5 complete defecation cycles, each one marked by a posterior body wall contraction (pBoc) (Iwasaki et al. 1995; Liu and Thomas 1994). The time interval between pBoc events was recorded as the DMP interval, and the number of expulsion (exp) events observed during the 5 recorded defecation cycles were noted. Animals were removed from the plate after recording to ensure that 10 unique individuals were assayed per strain. Data were plotted in Prism 9 and a one-way ANOVA was performed with a Bonferroni correction for multiple testing to calculate *P* values.

### Pharyngeal pumping assays

One-minute videos of 10 unique day 1 adult animals per genotype were recorded using a Nikon SMZ-18 Stereo microscope with a DS-Qi2 monochrome camera. Animals were permitted to freely move and feed on *E. coli* OP50 NGM plates during each recording. The number of pharyngeal pumps per minute were measured by reducing the video speed by 50% while counting. Data were plotted in Prism 9 and a one-way ANOVA was performed with a Bonferroni correction for multiple testing to calculate *P* values.

### Imaging and analysis of intracellular pH and intestinal lumen pH

For intracellular pH imaging, day 1 adult transgenic animals expressing pHluorin (Miesenböck et al. 1998) under the control of the pan-intestinal *nhx-2* promoter were used to visualize changes in intracellular pH, as previously described (Nehrke 2003).

The luminal pH measurements were obtained using Oregon Green, as previously described (Allman et al. 2009). Briefly, day 1 adults were washed from their plates and pelleted by brief centrifugation. The animals were resuspended in S-basal buffer with bacterial food to a final volume of 50μL, and an additional 50μL of S-basal containing 5mg/ml of an Oregon Green-dextran conjugate (70 kDa, Molecular Probes) was added. The worms were loaded with the conjugate by feeding for 2 hours at 20°C on a rocking platform and returned to NGM plates seeded with *E. coli* OP50 for imaging. The fluorescence excitation ratio was converted to pH using an *in vitro* calibration curve for the dye in solution.

For both assays, the plates were inverted onto the stage of a Nikon Eclipse TE2000-S microscope (Nikon Instruments), and freely moving subjects were kept within the field of view manually via stage manipulation. Illumination was provided by a Polychrome IV monochromator (TILL Photonics). Emissions were collected on a high-speed charge-coupled device camera (PCO Imaging). pH was assessed via dual excitation ratio spectroscopy, with sequential 10ms excitation at 410 and 470nm with 535nm emissions (pHluorin, intracellular pH) or sequential 10ms excitation at 440 and 490nm with 535nm emissions (Oregon Green-dextran, luminal pH). A 2×2 binning approach was utilized to increase signal to noise. Background subtraction was performed pixel-by-pixel at each excitation wavelength using images acquired from seeded plates lacking worms. Ratio images were further refined by thresholding, with the lowest intensity pixels (generally <2% of the data) set to zero. For quantitative analysis, fluorescence ratios were converted to pH using a Boltzmann equation and a calibration curve generated by *ex vivo* intestinal perfusion with a high K^+^/nigericin solution. TILLvisION software (TILL Photonics) was used for data analysis. Data were plotted using Prism 8.

### Raw data

The underlying numerical data and sample sizes for each individual experiment can be found in S1 Data.

## AUTHOR CONTRIBUTIONS

S.K.T., T.C., K.W.N., and R.H.D. designed the experiments. S.K.T., P.C.B., T.C., N.R.C., K.N.S., and K.R.V. performed the experiments. S.K.T., T.C., K.W.N., K.N.S., and R.H.D. interpreted the results. S.K.T. prepared the original draft of the manuscript. S.K.T., T.C., K.W.N., and R.H.D. reviewed and edited the manuscript. R.H.D. and K.W.N. secured funding for the study.

## ACKNOWLEDGEMENTS

The *Caenorhabditis* Genetics Center provided some of the strains used in this study, which is supported by the NIH Office of Research Infrastructure Programs (P40 OD010440).

