## Supplementary Material for "The TWK-26/KCNK3 potassium channel and FLR-4 protein kinase coordinate nutrient absorption in the *C. elegans* intestine"

| Strain | Genotype |
|---|---|
| N2 | Wild-type |
| DLS362 | *drl-1(rhd109) IV* |
| DLS523 | *mgIs70[*P*vit-3::GFP] I; flr-4(ut7) X* |
| DLS537 | *rhdSi42[*P*vit-3::mCherry::unc-54 3'UTR + cb-unc-119(+)] II* |
| DLS588 | *mgIs70[*P*vit-3::GFP] I; twk-26(rhd182[G485E]) flr-4(ut7) X* |
| DLS636 | *rhdSi42[*P*vit-3::mCherry::unc-54 3'UTR + cb-unc-119(+)] II; drl-1(rhd109) IV* |
| DLS638 | *rhdSi42[*P*vit-3::mCherry::unc-54 3'UTR + cb-unc-119(+)] II; flr-4(ut7) X* |
| DLS657 | *rhdSi42[*P*vit-3::mCherry::unc-54 3'UTR + cb-unc-119(+)] II; gpla-1(rhd117) V; flr-4(ut7) X* |
| DLS663 | *rhdSi42[*P*vit-3::mCherry::unc-54 3'UTR + cb-unc-119(+)] II; gpla-1(rhd117) V* |
| DLS664 | *gpla-1(rhd117) V* |
| DLS674 | *reSi5[*P*ges-1::TIR1::F2A::mTagBFP2::NLS::AID::tbb-2 3'UTR] I; rhdSi42[*P*vit-3::mCherry::unc-54 3'UTR + cb-unc-119(+)] II; flr-4(rhd209[mNG::TEV::3xFLAG::AID::flr-4]) X* |
| DLS719 | *rhdSi42[*P*vit-3::mCherry::unc-54 3'UTR + cb-unc-119(+)] II; twk-26(rhd182[G485E]) flr-4(ut7) X* |
| DLS735 | *rhdSi42[*P*vit-3::mCherry::unc-54 3'UTR + cb-unc-119(+)] II; twk-26(rhd182[G485E]) X* |
| DLS857 | *twk-26(rhd182[G485E]) X* |
| DLS865 | *twk-26(rhd182[G485E]) flr-4(ut7) X* |
| DLS870 | *reSi5[Pges-1::TIR1::F2A::mTagBFP2::NLS::AID::tbb-2 3'UTR] I; rhdSi42[*P*vit-3::mCherry::unc-54 3'UTR + cb-unc-119(+)] II; twk-26(rhd296[L232fs]) flr-4(rhd209[mNG::TEV::3xFLAG::AID::flr-4]) X* |
| DLS890 | *tir-1(ums63[tir-1::wrmScarlet]) III; twk-26(rhd182[G485E]) X* |
| DLS893 | *rhdSi42[*P*vit-3::mCherry::unc-54 3'UTR + cb-unc-119(+)] II; drl-1(rhd109) IV; twk-26(rhd182[G485E]) X* |
| DLS915 | *tir-1(ums63[tir-1::wrmScarlet]) III; twk-26(rhd182[G485E]) flr-4(ut7) X* |
| DLS942 | *rhdSi42[*P*vit-3::mCherry::unc-54 3'UTR + cb-unc-119(+)] II; gpla-1(rhd117) V; flr-1(ut11) X* |
| DLS962 | *tir-1(ums63[tir-1::wrmScarlet]) III; flr-4(ut7) X* |
| DLS964 | *twk-26(rhd182[G485E]) X; sqIs11[*P*lgg-1::mCherry::GFP::lgg-1 + rol-6(su1006)]* |
| DLS966 | *rhdSi42[*P*vit-3::mCherry::unc-54 3'UTR + cb-unc-119(+)] II; twk-26(rhd316[G485E, ΔS503-E519, T502fs]) X* |
| DLS975 | *twk-26(rhd182[G485E]) X; sqIs19[*P*hlh-30::HLH-30::GFP + rol-6(su1006)]* |
| DLS980 | *gpla-1(rhd117) V; flr-4(ut7) X* |
| DLS981 | *twk-26(rhd182[G485E]) flr-1(ut11) X* |
| DLS982 | *gpla-1(rhd117) V; flr-1(ut11) X* |
| DLS990 | *unc-119(tm4063)? III; twk-26(rhd182) X; wgIs396[hlh-11::TY1::EGFP::3xFLAG + unc-119(+)]* |
| DLS991 | *rhdSi42[*P*vit-3::mCherry::unc-54 3'UTR + cb-unc-119(+)] II; twk-26(rhd327[twk-26::2xminiIAA7::3xHA, G485E]) X* |
| DLS992 | *reSi7[*P*rgef-1::TIR1::F2A::mTagBFP2::NLS::AID::tbb-2 3UTR] I; rhdSi42[*P*vit-3::mCherry::unc-54 3'UTR + cb-unc-119(+)] II; twk-26(rhd327[twk-26::2xminiIAA7::3xHA, G485E]) X* |

| | |
|---|---|
| DLS994 | *reSi5[Pges-1::TIR1::F2A::mTagBFP2::NLS::AID::tbb-2 3'UTR] I; rhdSi42[*P*vit-3::mCherry::unc-54 3'UTR + cb-unc-119(+)] II; twk-26(rhd327[twk-26::2xminiIAA7::3xHA, G485E]) X* |
| DLS1020 | *pha-1(e2123)? III; twk-26(rhd182[G485E]) flr-4(ut7) X; rnyEx6[*P*nhx-2::pHluorin + pha-1(+)]* |
| DLS1021 | *pha-1(e2123)? III; flr-4(ut7) X; rnyEx6[*P*nhx-2::pHluorin + pha-1(+)]* |
| DLS1031 | *twk-26(rhd337[L229N]) X* |
| DLS1041 | *pha-1(e2123)? III; twk-26(rhd182[G485E]) X; rnyEx6[*P*nhx-2::pHluorin + pha-1(+)]* |
| DLS1042 | *twk-26(rhd343[twk-26::SL2::H2B::GFP]) X* |
| DLS1045 | *unc-119(ed3) III; rhdEx109[*P*twk-26::mCherry* + P*myo-3::GFP + unc-119(+)]* |
| JC51 | *flr-4(ut7) X* |
| JC55 | *flr-1(ut11) X* |
| KWN26 | *pha-1(e2123) III; rnyEx6[*P*nhx-2::pHluorin + pha-1(+)]* |
| MAH215 | *sqIs11[*P*lgg-1::mCherry::GFP::LGG-1 + rol-6(su1006)]* |
| MAH235 | *sqIs19[*P*hlh-30::HLH-30::GFP + rol-6(su1006)]* |
| OP396 | *unc-119(tm4063) III; wgIs396[hlh-11::TY1::eGFP::3xFLAG + unc-119(+)]* |
| RPW403 | *tir-1(ums63[tir-1::wrmScarlet]) III* |

**Supplementary Table 1. The *C. elegans* strains used in this study.**

| **Target Gene** | **Location in gene, crRNA guide number** | **crRNA sequence** | **Alleles** | **Genomic edit** |
|---|---|---|---|---|
| *twk-26* | middle, rhd56 | CUAGUAGUGCUUGGUGACAU | *rhd296* | L232fs |
| *twk-26* | C-terminus, rhd43 and rhd54 | AGGAGAGUGAUGUGGAGUGA | *rhd316* | ΔS503-E519 |
| *twk-26* | C-terminus, rhd43 | AGGAGAGUGAUGUGGAGUGA | *rhd327* | twk-26:: 2xminiIAA7::3xHA |
| *twk-26* | middle, rhd56 | CUAGUAGUGCUUGGUGACAU | *rhd337* | L229N |
| *twk-26* | C-terminus, rhd43 | AGGAGAGUGAUGUGGAGUGA | *rhd343* | twk-26:: SL2::H2B::GFP |

**Supplementary Table 2. The crRNAs used in this study.**

| mRNA Target | Primer Sequence (5' to 3') |
|---|---|
| *nlp-40* (amplicon #1) | **Forward:** CCGACACATTCCTTGGATTTG<br>**Reverse:** CTCCGGCTCTCAACACTTC |
| *nlp-40* (amplicon #2) | **Forward:** ATTCTGCTATCTTTTGTTGCGAC<br>**Reverse:** GGTTGCTCCTTTTGCTTCAC |
| *nhx-2* (amplicon #1) | **Forward:** CTTTGTGGAGAACTTTTCGGTC<br>**Reverse:** GATTCTGTGGCTTTTACCGTG |
| *nhx-2* (amplicon #2) | **Forward:** TGCTTATTTTGAATGGGAGAATGTC<br>**Reverse:** CGGGAAAAGTTCATTGAGATGTG |
| *pept-1* (amplicon #1) | **Forward:** TCAGATCTCAACCATGCCTTG<br>**Reverse:** TGAACCTCCCATAAATACCAGTG |
| *pept-1* (amplicon #2) | **Forward:** GAGAGTTCTTCCCCATGACAAG<br>**Reverse:** ATATGTGAGAGTCCAATCGCC |

**Supplementary Table 3. The RT-qPCR primers used in this study.**

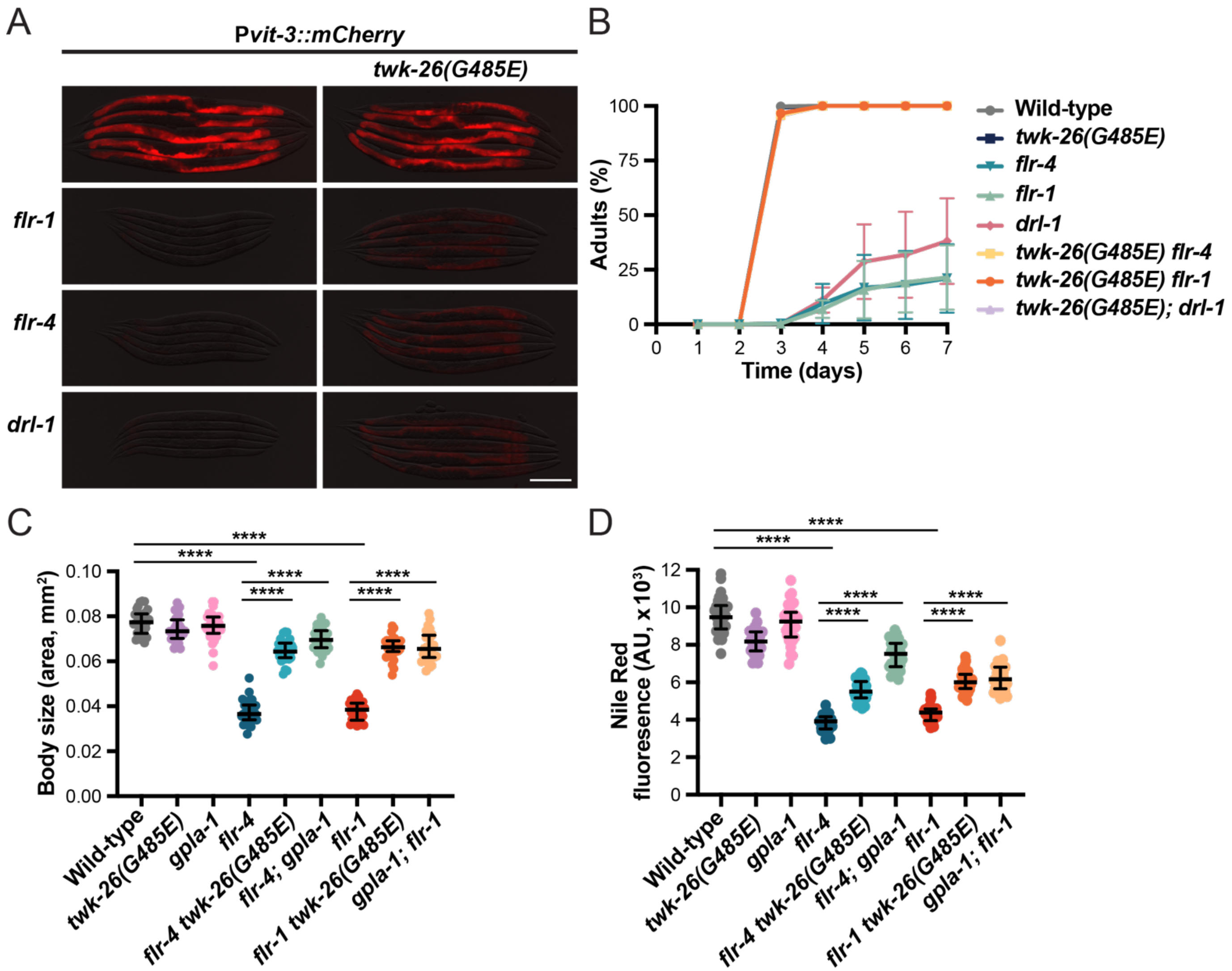


**Figure S1. A mutation in *twk-26* suppresses the growth and lipid metabolism defects caused by loss of DRL-1, FLR-4, or FLR-1.** (A) Representative overlaid DIC and P*vit-3::mCherry* fluorescence images of day 1 adult wild-type or mutant animals (scale bar, 200μm). (B) Growth rate of the indicated strains reared from embryos at 20°C. Data are shown as the mean +/- SEM of 3 independent experiments. Some of these data are also shown in Figure 1E. (C) Body size of fixed (60% isopropanol) day 1 adult animals of the indicated genotypes (median and interquartile range; ns, not significant, **, *P*=0.0012, ****, *P*<0.0001, one-way ANOVA). (D) Quantification of Nile Red staining of wild-type and mutant day 1 adults (median and interquartile range; ns, not significant, ****, *P*<0.0001, one-way ANOVA). (A-D) All strains contain *rhdSi42*.

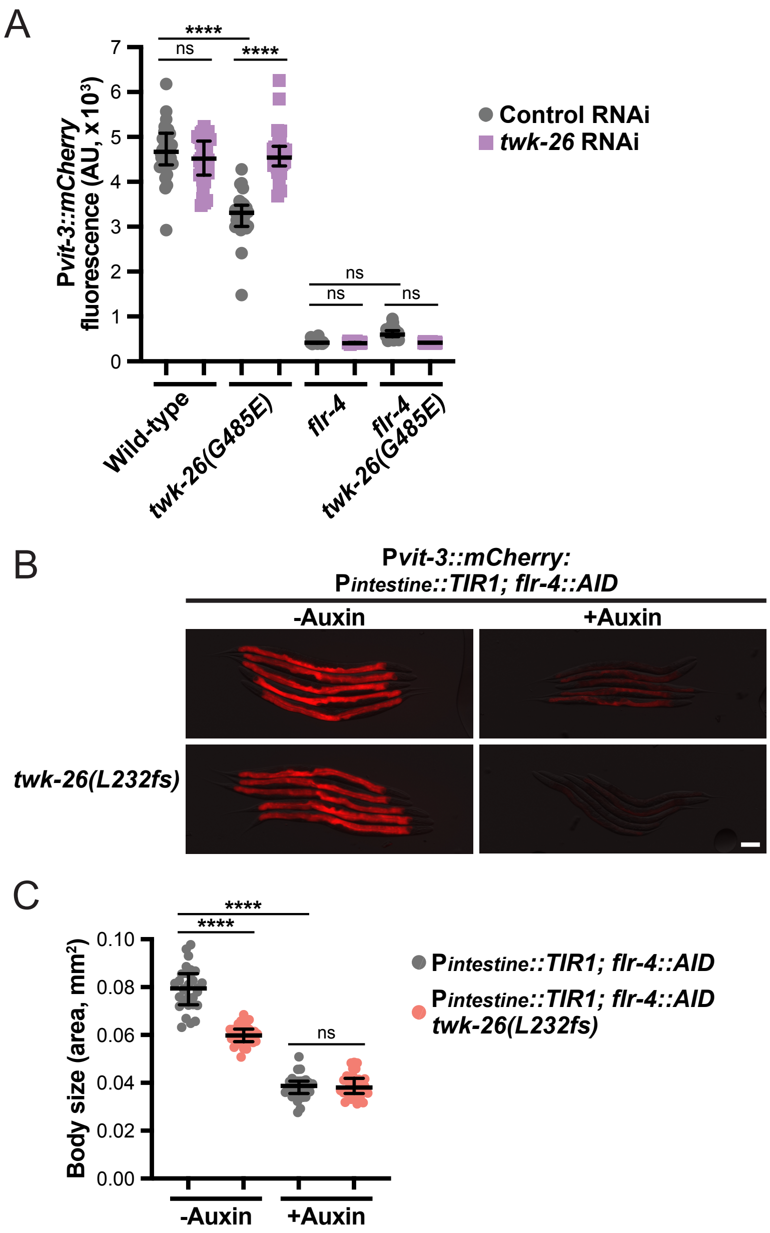


**Figure S2. Loss of *twk-26* fails to suppress the *flr-4* mutant phenotypes.** (A) Quantification of P*vit-3::mCherry* fluorescence in day 1 adult animals subjected to control or *twk-26* RNAi (median and interquartile range; ns, not significant, ****, *P*<0.0001). (B) Representative overlaid DIC and mCherry fluorescence images of AID::FLR-4 day 1 adult animals expressing P*ges-1::TIR1* (intestinal driver of TIR1) in the absence or presence of 4mM auxin (scale bar, 100µm). (C) Body size measurements of day 1 adult AID::FLR-4 animals expressing P*ges-1::TIR1* in the absence or presence of 4mM auxin. Data are displayed as the median and interquartile range (ns, not significant, *P*<0.0001, one-way ANOVA). The strains all contain the *rhdSi42* transgene.

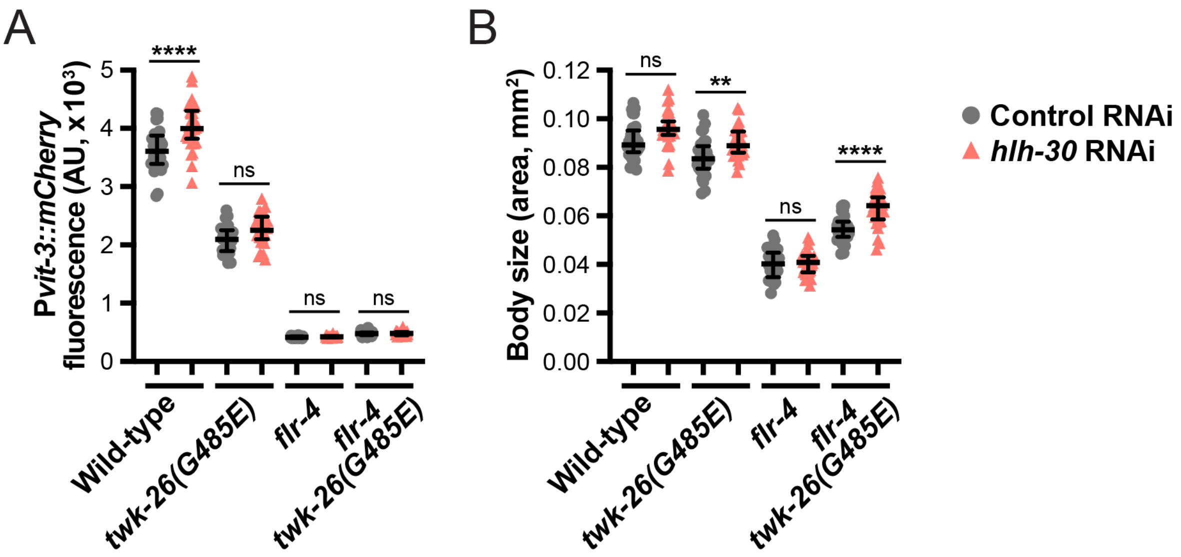


**Figure S3. Genetic suppression of the *flr-4* mutation by TWK-26 G485E does not require *hlh-30*.** Quantification of (A) body size and (B) P*vit-3::mCherry* fluorescence in day 1 adult animals subjected to control or *hlh-30* RNAi. Data are presented as median and interquartile range (ns, not significant, **, *P*=0.0072, ****, *P*<0.0001, one-way ANOVA). All strains contain *rhdSi42*.

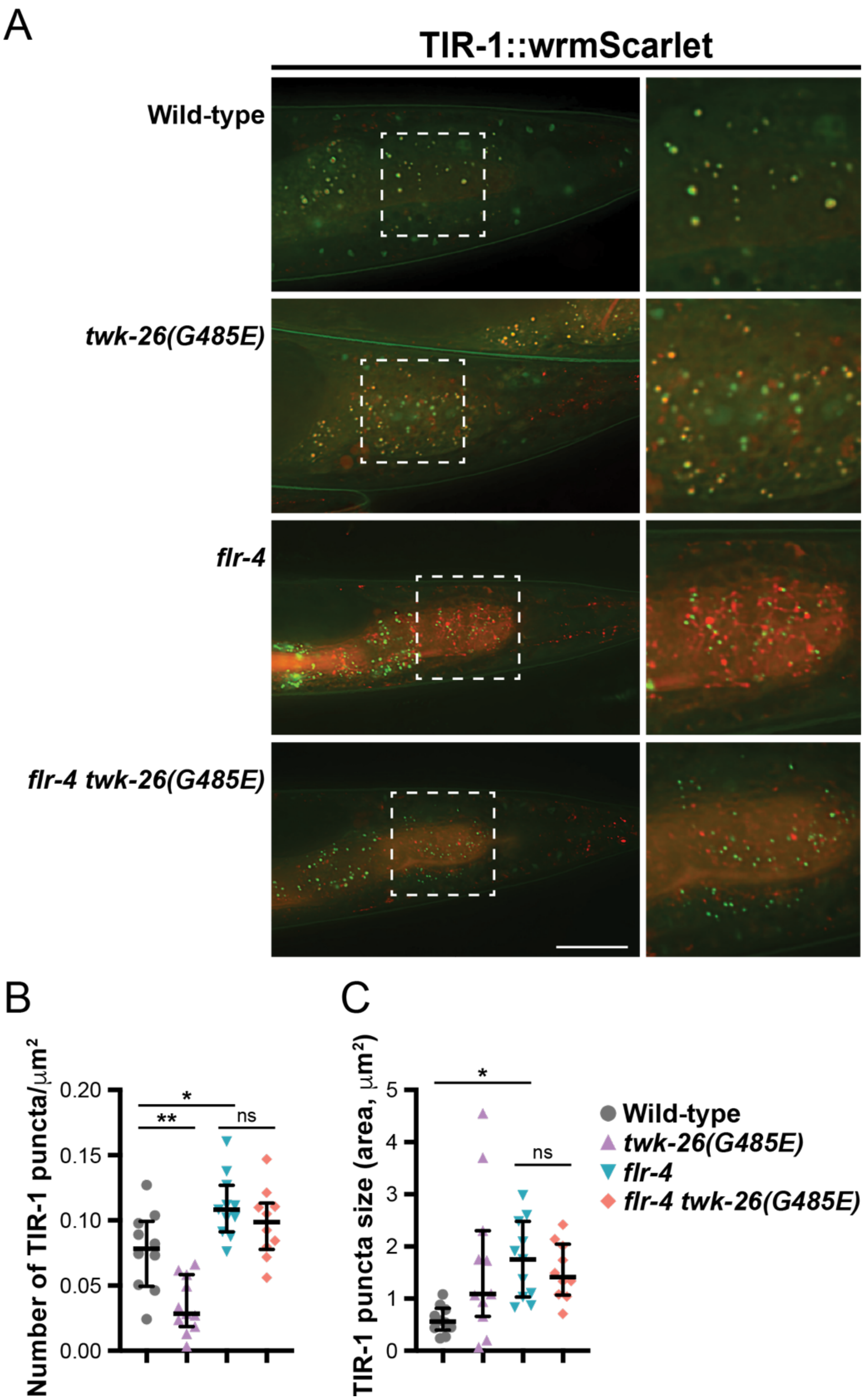


**Figure S4. The *twk-26(G485E)* mutation does not suppress hyperactivation of the p38/PMK-1 pathway in the *flr-4* mutant.** (A) Representative fluorescent images, and magnified insets (right panels), of TIR-1::wrmScarlet puncta in posterior intestinal cells of day 1 adult wild-type and mutant animals (scale bar, 25µm). Yellow puncta are autofluorescent gut granules, which fluoresce in both the green and red channels. Quantification of (B) the total number and (C) the average size of TIR-1 puncta in the posterior intestine (median and interquartile range, ns, not significant, *, *P*<0.05. **, *P*<0.01, ****, *P*<0.0001, one-way ANOVA).

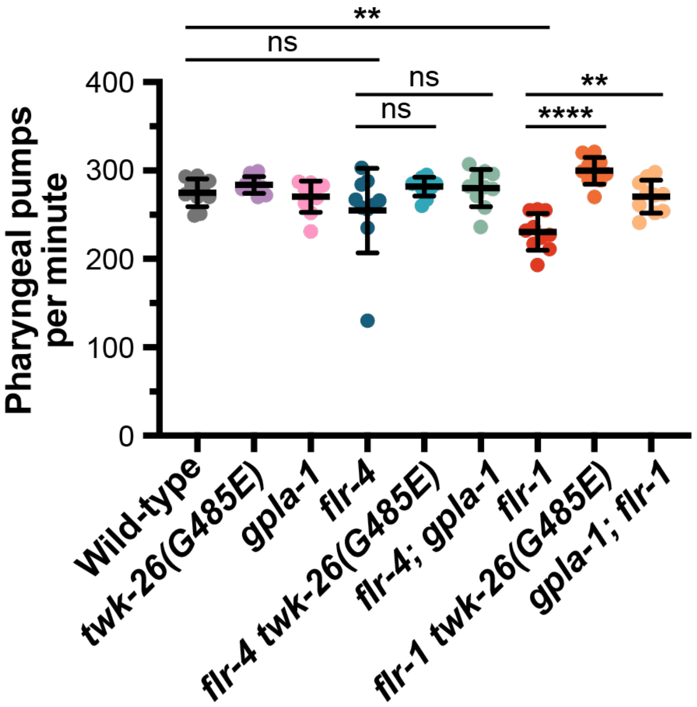


**Figure S5. Loss of *flr-4* does not impair pharyngeal pumping.** Quantification of pharyngeal pumping rates (number of pumps per minute) of 10 individuals per genotype. Data are presented as the mean +/- SD (ns, not significant, **, *P*<0.01, ****, *P*<0.0001, one-way ANOVA).

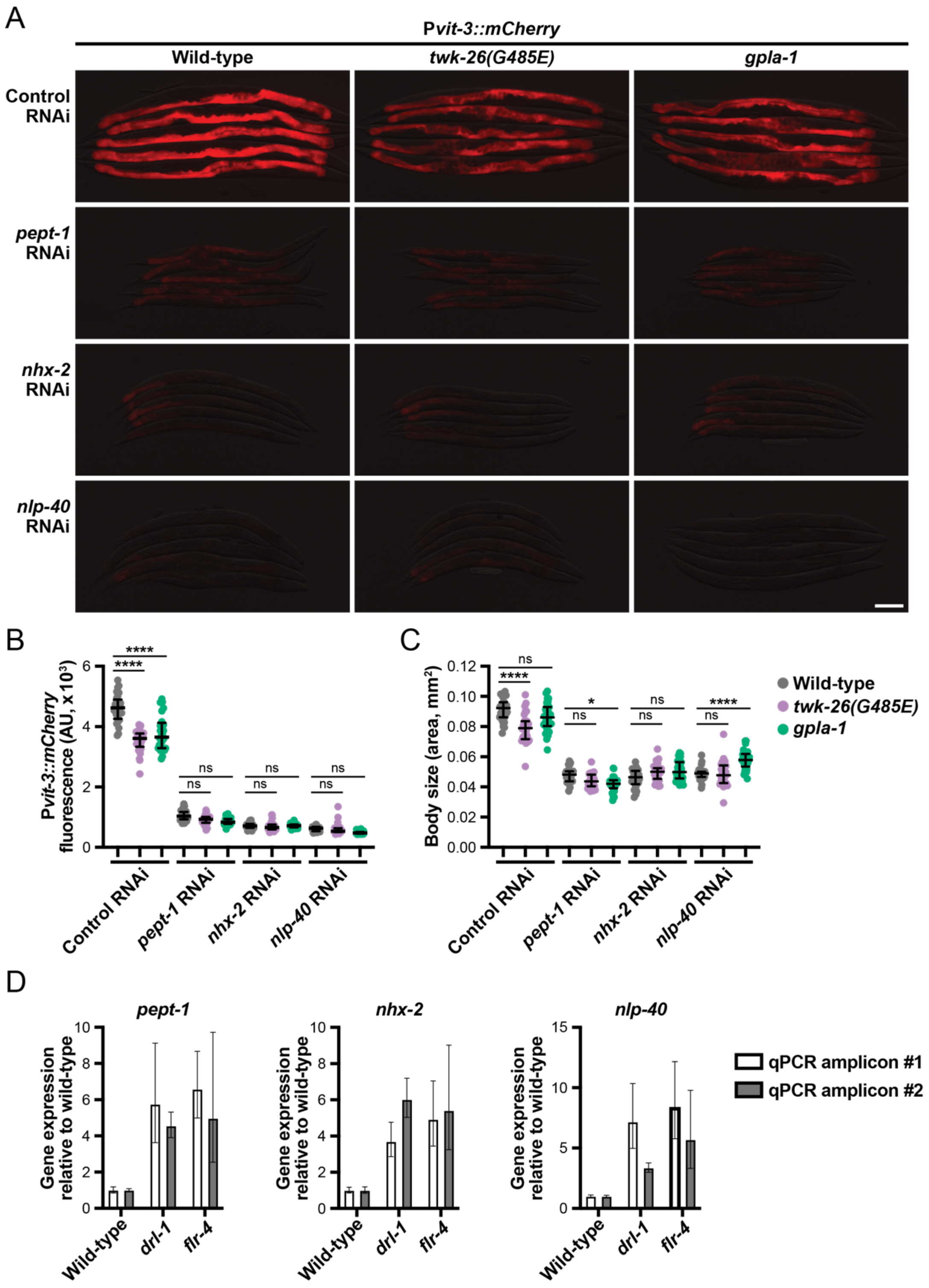


**Figure S6. Genes that function in dipeptide transport and the DMP are required for metabolic homeostasis but function independently of *twk-26*.** (A) Representative overlaid DIC and mCherry fluorescence images of day 1 adult wild-type, *twk-26(G485E),* and *gpla-1* animals subjected to the indicated RNAi treatment (scale bar, 200μm). Quantification of (B) P*vit-3::mCherry* fluorescence and (C) body size of day 1 adult wild-type, *twk-26(G485E),* and *gpla-1* animals following knock-down of the indicated genes by RNAi (median and interquartile range; ns, not significant, *, $P$=0.0374, ****, $P$<0.0001, one-way ANOVA). (A-C) Animals contain the *rhdSi42* transgene. (D) Expression of the indicated genes in day 1 adult animals, as measured by RT-qPCR (two qPCR amplicons per gene).

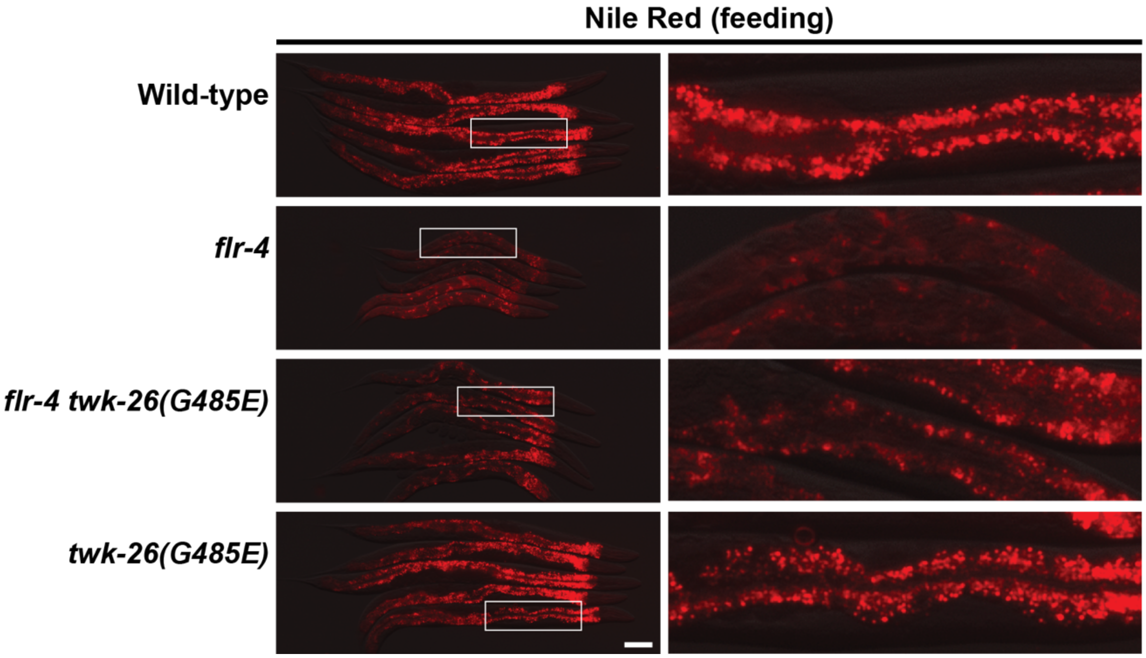


**Figure S7. TWK-26 G485E restores Nile Red absorption in the *flr-4* mutant.** Representative fluorescence images, and magnified insets (right panels), of day 1 adult animals fed *E. coli* OP50 supplemented with Nile Red (scale bar, 100μm).

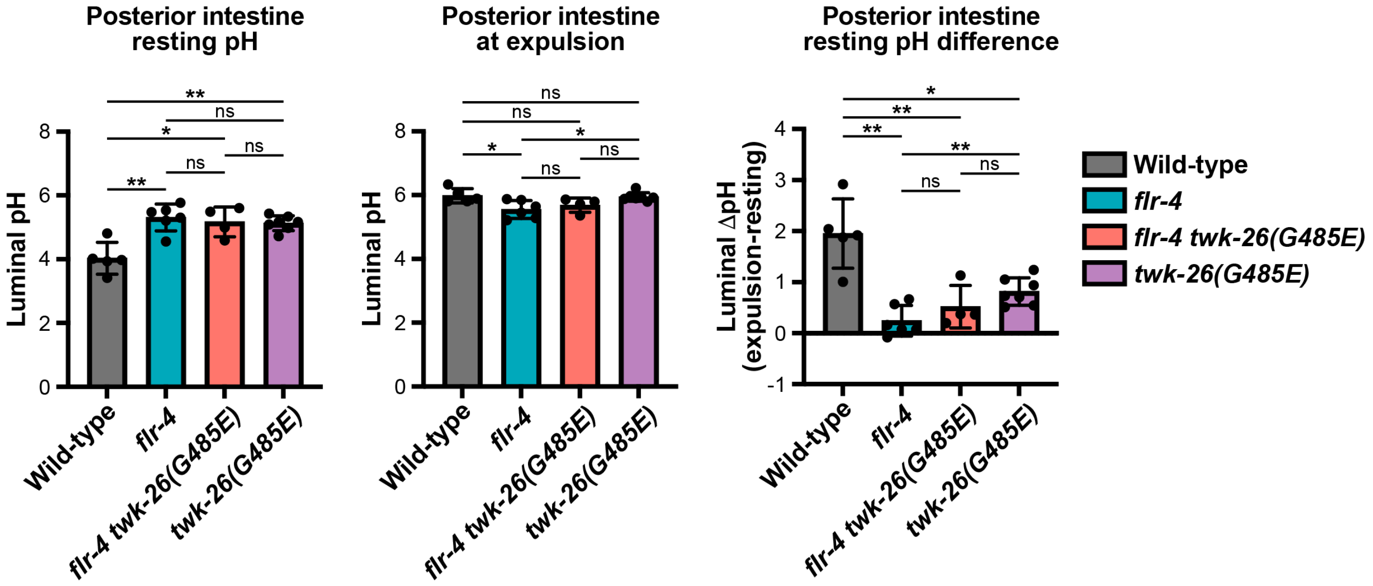


**Figure S8. The *flr-4* mutant fails to alkalinize its posterior intestine following defecation.** Measurements of posterior luminal pH during rest (left panel) and following pBoc/expulsion (middle panel) in animals fed dextran-conjugated Oregon Green 488. The difference between the resting and expulsion luminal pH is also reported (right panel). Data are presented as mean +/- SD (N=4-7 animals per genotype; ns, not significant, *, $P<0.05$, **, $P<0.01$, Brown-Forsythe and Welch ANOVA followed by Fishers LSD Test).
